# Divergent iron-regulatory states contribute to heterogeneity in breast cancer aggressiveness

**DOI:** 10.1101/2023.06.23.546216

**Authors:** William D. Leineweber, Maya Z. Rowell, Sural Ranamukhaarachchi, Alyssa Walker, Yajuan Li, Jorge Villazon, Aida Mestre Farrera, Zhimin Hu, Jing Yang, Lingyan Shi, Stephanie I. Fraley

## Abstract

Primary tumors with similar mutational profiles can progress to vastly different outcomes where transcriptional state, rather than mutational profile, predicts prognosis. A key challenge is to understand how distinct tumor cell states are induced and maintained. In triple negative breast cancer cells, invasive behaviors and aggressive transcriptional signatures linked to poor patient prognosis can emerge in response to contact with collagen type I. Herein, collagen-induced migration heterogeneity within a TNBC cell line was leveraged to identify transcriptional programs associated with invasive versus non-invasive phenotypes and implicate molecular switches. Phenotype-guided sequencing revealed that invasive cells upregulate iron uptake and utilization machinery, anapleurotic TCA cycle genes, actin polymerization promoters, and a distinct signature of Rho GTPase activity and contractility regulating genes. The non-invasive cell state is characterized by actin and iron sequestration modules along with glycolysis gene expression. These unique tumor cell states are evident in patient tumors and predict divergent outcomes for TNBC patients. Glucose tracing confirmed that non-invasive cells are more glycolytic than invasive cells, and functional studies in cell lines and PDO models demonstrated a causal relationship between phenotype and metabolic state. Mechanistically, the OXPHOS dependent invasive state resulted from transient HO-1 upregulation triggered by contact with dense collagen that reduced heme levels and mitochondrial chelatable iron levels. This induced expression of low cytoplasmic iron response genes regulated by ACO1/IRP1. Knockdown or inhibition of HO-1, ACO1/IRP1, MRCK, or OXPHOS abrogated invasion. These findings support an emerging theory that heme and iron flux serve as important regulators of TNBC aggressiveness.

## Introduction

Metastasis causes 60-90% of cancer deaths (1, 2), and many experts believe that targeting initial metastatic invasion and secondary spread will be the most effective strategy for reducing cancer mortality (2, 3). Despite this clinical impetus, there are few effective treatments that specifically target metastasis (4). This is in part because the genetic determinants of tumor initiation and early progression do not distinguish whether a tumor will metastasize (5–8). Instead, many cancers acquire metastatic capabilities by co-opting programs necessary for normal physiological processes (9–12). It is particularly unclear how cancer cells in the primary tumor initiate and sustain transcriptional programs that would normally be transient under physiological conditions to carry out metastatic functions. Accumulating evidence suggests that exposure to various microenvironmental factors could induce stable transcriptional responses that enable metastatic capabilities (13, 14). However, questions remain regarding why only a subset of cells exposed to such factors acquire aggressive phenotypes, and what features define these aggressive phenotypes (15).

Recently, collective cell invasion has emerged as a new paradigm defining an important initial step within the overall metastatic cascade (16). Aggressive solid tumors across multiple cancer types display strands of collectively invasive tumor cells (17, 18) that have been linked to circulating tumor cell clusters and have greater metastatic potential than single cells (19). Understanding why and how some cancer cells adopt a collectively invasive phenotype in the primary tumor microenvironment could lend insight into valuable therapeutic targets. *In vitro* models of collective invasion induce this phenotype by embedding cells in dense extracellular matrices (ECMs) (20–22). Dense ECM is a hallmark of the tumor microenvironment (TME) and contributes both to tumorigenesis and metastasis, especially in breast cancer patients (23–26). Bulk sequencing analysis has shown that cancer cells upregulate a transcriptional program that is predictive of poor prognosis when embedded in dense collagen type I prior to undergoing collective invasion, and this response is related to oxidative stress (21, 27).

Here, we identify the mechanisms driving and sustaining the collective invasion phenotype of triple negative breast cancer (TNBC) cells in response to dense type I collagen. Leveraging the fact that not all cells within a population respond in the same way, we analyze single cell sequencing of collectively invasive versus non-invasive migration phenotypes. We find that these distinct phenotypes are associated with divergent states of iron, cytoskeletal, and metabolism regulation. Mechanistically, our experiments show that dense Col1 induces HO-1 expression and activity, which catabolizes heme and releases iron, carbon monoxide (CO), and biliverdin. This leads to two subpopulations of cells. The minority subpopulation is non-invasive, displays higher levels of mitochondrial labile iron, and expresses genes associated with high cytoplasmic iron that promote iron storage. This echoes the canonical response to HO-1 activity in normal cells. The majority subpopulation is invasive and displays reduced mitochondrial labile iron, which is associated with mitochondrial biosynthetic activity. This population also upregulates heme transport and ISC containing genes as well as genes associated with an IRP-mediated response to low cytoplasmic iron that promotes iron uptake, including the actin-myosin regulatory kinase MRCK. We show evidence for these states in human breast tumors and find that they are predictive of opposing clinical outcomes. Knockdown of HO-1 or IRP1 or inhibition of MRCK abrogates the invasive phenotype associated with a low cytoplasmic iron response and high mitochondrial iron utilization. The invasive state is also more dependent on OXPHOS while the non-invasive is more glycolytic. Inhibition of glycolysis or OXPHOS targets these distinct populations in high density (HD) Col1 and in a collagen-rich PDO model, demonstrating the relevance of this ECM-driven mechanism of metabolic and phenotypic heterogeneity in TNBC. Overall, this study suggests that dense collagen type I can drive metabolic, cytoskeletal, and phenotypic heterogeneity in TNBC through HO-1 activation and its effects on heme and iron pools. This provides a systems level model for the divergent role that collagen, HO-1, and iron can play in TNBC and implicates specific biomarkers and therapeutic strategies for further investigation.

## Results

### Distinct modes of cytoskeletal regulation underlie TNBC migration heterogeneity, with MRCK necessary for collective invasion.

Previously, we developed a method to investigate phenotypic heterogeneity within individual cell types. Cells expressing a photoconvertible protein construct were “laser tagged” and sorted based on their dynamic characteristics for subsequent molecular and functional analyses. We demonstrated the utility of this method by sorting MDA-MB-231 (MDA) cells, a human TNBC cell line, and 4T1 cells, a mouse TNBC cell line, according to their migration phenotype in high density collagen I (Col1). When embedded as single cells at low seeding density, the majority of these cells undergo proliferation and collective invasion to form clonal network-like structures over the course of seven days. However, a small proportion form clonal non-invasive structures (Fig. 1A, B) (21, 22, 27, 28). Phenotypic sorting followed by single cell RNA sequencing (scRNAseq) at 7 days of culture (Fig. 1 C, Table S1) generated a rich dataset (28, 29). Here, we set out to conduct a deeper analysis of this dataset with the goal of identifying the mechanisms that initiate and sustain the two emergent migration phenotypes within a TNBC cell population.

**Figure 1:**
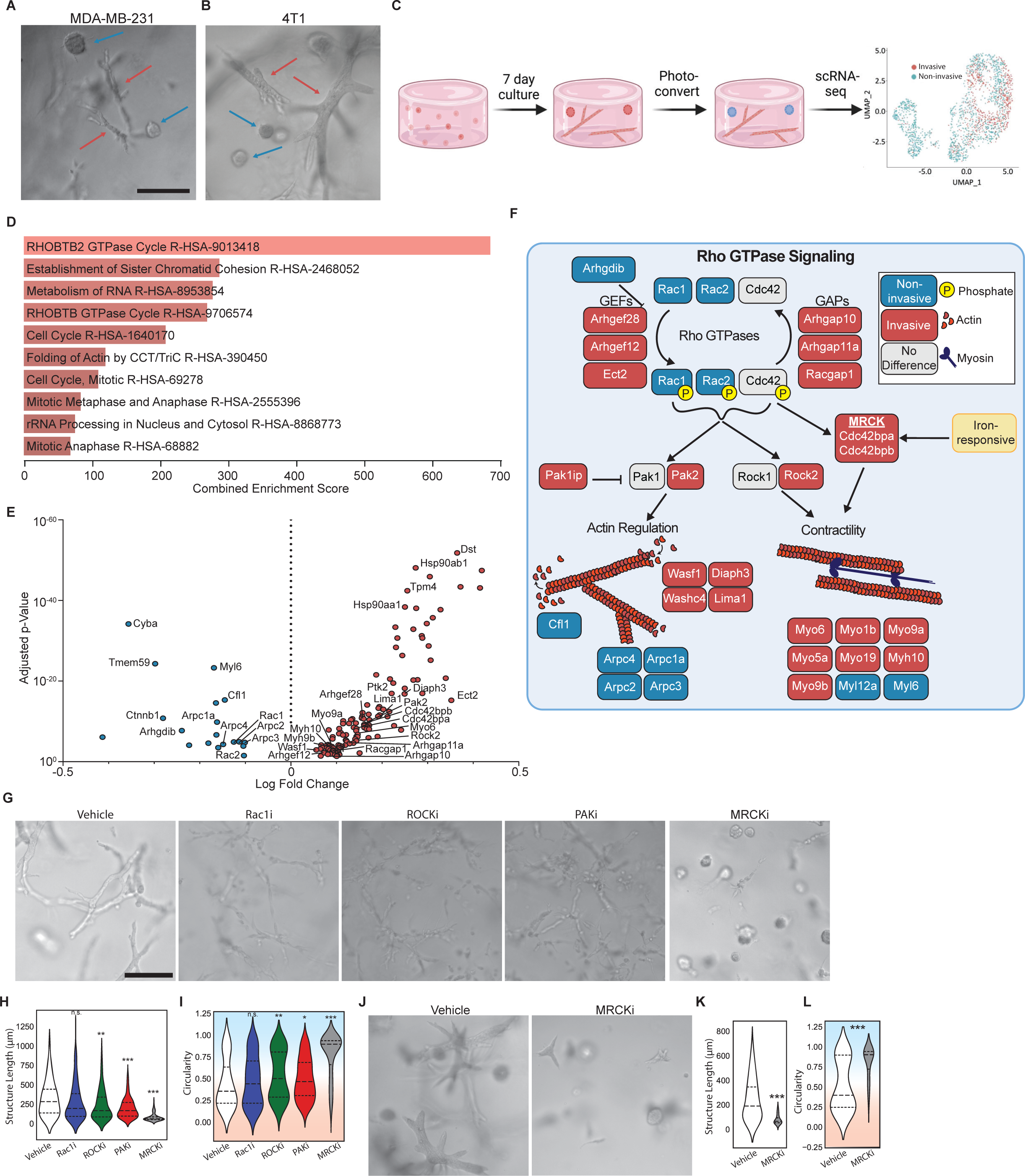
Distinct cytoskeletal programs underlie breast cancer migration heterogeneity, with MRCK serving as a critical regulatory node of collective invasion. **(A-B)** Representative micrographs of MDA-MB-231 (A) and 4T1 (B) collective migration phenotypes that form when single cell suspensions embedded in 6mg/ml collagen I are cultured for one week. Red arrows point to the invasive phenotype, while the blue arrows show the non-invasive spheroidal structures. **(C)** Schematic of phenotypically-supervised single cell RNA sequencing (PhenoSeq). Photoconversion of cells from invasive and non-invasive collective migration phenotypes enables subsequent sorting and sequencing while retaining knowledge of the cell’s phenotype. This labeling enables clustering based on phenotype, which reveals distinct gene expression patterns that algorithm-based unsupervised clustering misses. **(D)** Pathway enrichment analysis using the Reactome 2022 database of all differentially expressed genes found in the MDA-MB-231 PhenoSeq data. **(E)**Volcano plot of differential gene expression between cells in the collective non-invasive and invasive phenotypes related to Rho GTPase signaling. **(F)** Diagram showing Rho GTPase signaling and the downstream effects on actin cytoskeleton regulation and actomyosin contractility. The MRCKα-encoding gene *Cdc42bpa* contains an iron responsive element in the 3’ UTR of the mRNA. **(G)** Representative micrographs showing how cytoskeletal inhibitors alter the collective migration phenotype of MDAs within Col1 matrices. **(H-I)** Quantification of the multicellular structures showing length (H) and the circularity (I) of the structures. **(J)** Representative micrographs of 4T1 multicellular structures after seven days in Col1 matrices with vehicle or MRCK inhibitor treatment. **(K-L)** Quantification of the phenotypes showing the length (Feret Diameter) (N) and circularity (O) of the structures. Background coloring is used as a visual aid to show that high circularity values are linked to the non-invasive phenotype (blue background), while the more invasive phenotypes have lower circularity (red background). Scale bar = 200μm. N = 3 biological replicates per treatment group. Horizontal lines within the violin plots show the quartiles of the data. Statistical significance was determined using Student’s t-test or one-way ANOVA followed by Tukey post-tests. * p < 0.05, ** p < 0.01, *** p < 0.001, **** p < 0.0001.

First we analyzed expression differences in genes that comprise the cell motility machinery, with the hypothesis that this would reveal key components that sustain invasive versus non-invasive behaviors. Pathway enrichment analysis of genes that were differentially expressed (DE) between the invasive and non-invasive cells showed that several migration related pathways were among the most highly enriched, including annotations for RHOBTB2 GTPase Cycle (combined score = 683.14, adjusted p-value = 4.319e-11) and Folding of Actin by CCT/TriC (combined score = 318.68, adjusted p-value = 2.93e-6) (Fig. 1 D). The phenotype-associated patterns of DE genes in these categories and additional gene ontologies (GO) annotated for Rho GTPase signaling suggested that distinct cytoskeletal programs may be active in invasive versus non-invasive cells (Fig. 1 E, Table S2). We mapped several of the most well-characterized genes onto a schematic of known Rho GTPase signaling and downstream effectors of actin cytoskeleton regulation and actomyosin contractility to illustrate this point (Fig. 1 F). Manually curated genes (*Pak1ip, Rac1, Washc4, Lima1, Myo5a, Myo19, Myo1b, Myl12a, Cfl1*) that were not included in the GO annotation, likely due to the bias towards more common genes (30, 31), are also included to further show implications for actin and contractility.

Invasive cells showed higher expression of genes encoding multiple guanine nucleotide exchange factors (GEFs) and GTPase-activating proteins (GAPs), which would be expected to promote more dynamic Rho GTPase signaling. Downstream cytoskeletal regulators expressed by invasive cells (*Diaph3, Lima1, Washc4, Wasf1, Pak2*) would be expected to promote actin polymerization and result in cell elongation (32–34). Additionally, actomyosin contractility genes upregulated in the invasive cells (*Rock2, Cdc42bpa & Cdc42bpb, Myo6, Myo1b, Myo9a, Myo5a, Myo9b, Myh10*) would be expected to enhance motility by enabling cells to generate traction and migrate through the ECM (35–37). *Myo6, Myo9a,* and *Myo9b* have previously been implicated in promoting collective migration (38–40).

Conversely, *Cfl1,* which promotes actin depolymerization and severing (41), and *Arhgdib*, which inhibits the dissociation of GDP and the subsequent binding of GTP to Rho proteins (42, 43), were more highly expressed in non-invasive cells. Counterintuitively, expression of *Rac1* and *Rac2* was upregulated in the non-invasive phenotype as well. Overall, this analysis suggested that numerous cytoskeletal regulators may act in concert with numerous myosin genes to functionally achieve the two TNBC cell migration phenotypes.

To begin to test whether these patterns of cytoskeletal gene expression are functional at the protein level, we selected well-characterized inhibitors targeting a subset of the DE genes. Inhibition of RAC1 had minimal effects on the prevalence of each phenotype or extent of collective invasion, though it did promote less connected invasive structures that appeared like beads-on-a-string (Fig. 1G). Immunostaining for RAC1 showed that expression was more heterogeneous within invasive structures, with some cells lacking expression, which may account for the overall lower expression of RAC1 in invasive structures (Fig. S1). Inhibition of ROCK and PAK also had minimal effects on the prevalence of each phenotype or extent of collective invasion, but promoted less connected invasive structures (Fig. 1G). In contrast, inhibition of MRCK (containing subunits *Cdc42bpa* & *Cdc42bpb)* produced a dramatic shift to the non-invasive phenotype. The length of the MRCK-inhibited (MRCKi) structures was 73.43 ± 50.28 μm (mean ± standard deviation), which was 4.83 ± 0.11 fold less than the vehicle (Fig. 1 H). The circularity of structures was 0.91 ± 0.08, reflecting the homogenization of the phenotypes to small spheroids (Fig. 1 I). Additional measurements of the phenotypes, such as the area, perimeter, minor Feret Diameter, roundness, and solidity are available in Fig. S2 A-E.

These phenotypic effects were observed at concentrations that do not significantly alter cell viability compared to the control (Fig. S2 F), and the MRCK inhibitor showed a strong dose-dependent response even as low as 10nM (Fig. S2 G-M). The shift to the non-invasive phenotype induced by MRCK was confirmed in 4T1 cells as well (Fig. 1 J-L, Fig. S2 N-R). These results suggested that the upregulation of the actin-myosin regulatory kinase MRCK plays an important role in enabling the invasiveness of TNBC cells. This prompted us to investigate regulators of MRCK expression.

### Distinct iron regulatory programs differentiate non-invasive from invasive TNBC cells

MRCK has been reported to colocalize with transferrin receptor 1 (*Tfrc*) and its subunit *Cdc42bpa* contains an iron response element in the 3’ UTR of its mRNA (44, 45). This led us to examine potential links between the invasive cytoskeletal regulatory program and iron regulation. A literature search did not reveal any direct links between the other DE actomyosin contractility genes and *Tfrc*. However, DE genes linked to iron were identified through pathway enrichment (Iron Uptake and Transport, combined score = 11.27, adjusted p-value = 0.036), iron-related genesets from GSEA, and manual curation of recently identified iron-related genes (Fig. 2 A). Invasive cells upregulated genes involved in iron uptake, including *Tfrc*, *Slc39a14,* and *Aco1*, while non-invasive cells upregulated genes involved in iron sequestration, such as *Fth1* and *Tmsb4x* (46). The strong enrichment of *Tfrc* in the invasive cells prompted us to also assess genes whose expression has previously been shown to be down-regulated by *Tfrc* knockout *(47)*. We found that the invasive phenotype is enriched for genes whose expression depends on *Tfrc* (Fig. 2 B).

**Figure 2:**
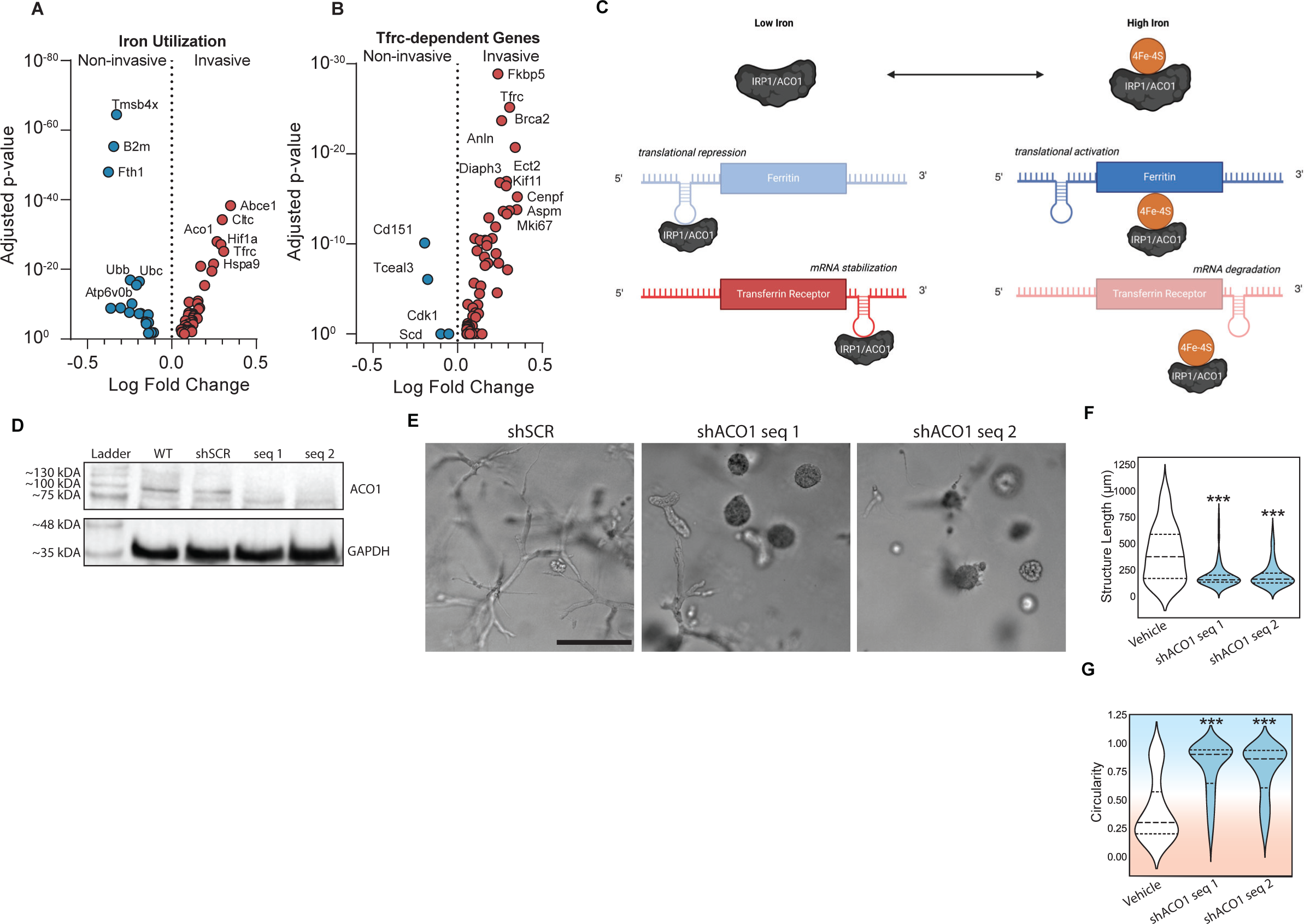
Divergent iron metabolism responses mediated by ACO1 differentiate non-invasive from invasive phenotypes. **(A)** Volcano plot of differentially expressed genes identified from PhenoSeq that are related to intracellular iron usage. **(B)** Volcano plot of differentially expressed genes whose expression decreases when *Tfrc* is knocked down. **(C)** Schematic of cytoplasmic iron regulation of transcription mediated by IRP1/ACO1. Adapted from (108) **(D)** Western blot confirming the knockdown in protein expression of ACO1 in MDAs following lentiviral transduction with two separate shRNA sequences, as compared to wild type (WT) and shRNA targeting a scramble control sequence (shSCR). **(E)** Representative micrographs of lentivirally-transduced MDAs after seven days in Col1 matrices. **(F-G)** Quantification shows the maximum length of structures (Feret Diameter) (Q) and circularity (R). Background coloring is used as a visual aid to show that high circularity values are linked to the non-invasive phenotype (blue background), while the more invasive phenotypes have lower circularity (red background). Scale bars = 200μm. N = 3 biological replicates per treatment group. Statistical significance was determined using one-way ANOVA followed by a Dunnett post-test comparing each treatment group to the vehicle group. * p < 0.05, ** p < 0.01, *** p < 0.001, **** p < 0.0001.

In normal cells, expression of *Cdc42bpa, Tfrc*, *Slc39a14, and Fth1* is regulated by cytoplasmic iron levels through *Aco1* (48). When cytoplasmic iron levels are low, ACO1 functions as iron response protein 1 (IRP1), binding to iron response elements (IREs) in the mRNAs of these genes (Fig. 2 C). Binding to *Cdc42bpa, Tfrc*, and *Slc39a14* mRNA promotes their expression, while binding to *Fth1* mRNA represseses its expression. When cytoplasmic iron levels are abundant, IRP1 is bound by an iron-sulfur cluster (ISC) and functions as an aconitase, *Aco1*, no longer binding to IREs in mRNAs (Fig. 2 C). This reverses the transcriptional pattern. The invasive transcriptional program is therefore consistent with IRP1-mRNA interactions in response to a low cytoplasmic iron state while the non-invasive program is consistent with a lack of IRP1-mRNA interactions mediated by a high cytoplasmic iron state. To test this hypothesis, MDAs were lentivirally transduced with shRNA targeting *Aco1/Irp1* or scramble control (shSCR), which was validated via western blot (Fig. 2 D). Knocking down IRP1/ACO1 significantly reduced invasion (Fig. 2 E-G, Fig. S3), suggesting that the invasive phenotype is indeed dependent on IRP1/ACO1.

### Iron-associated transcriptional states induced by HD Col1 are evident in TNBC patient tumors

To assess the relevance of these collagen-induced TNBC cell states to human breast cancer and clinical outcomes, we conducted survival curve analysis using the breast cancer RNA-seq dataset from the Km Plotter tool (49). The mean expression of the top iron-related genes from the non-invasive phenotype (*Tmsb4x, B2m, Fth1, Ubb, Ubc, Atp6v0b*) are associated with improved overall survival (HR = 0.32) in TNBC patients (Fig. 3 A). On the other hand, the top genes from the invasive phenotype (*Abce1, Ctlc, Aco1, Hif1a, Tfrc, Hspa9*) together predict worse survival (HR = 3.03) (Fig. 3 B). Analyzing these genes individually, we found that *Aco1* is significantly correlated with worse overall patient survival in all breast cancer patients (HR = 1.56, p = 0.00014) (Fig. 3 C), and TNBC patients had even worse outcomes when stratified by *Aco1* expression (Fig. 3 D) (HR = 4.22, p = 0.0041).

**Figure 3:**
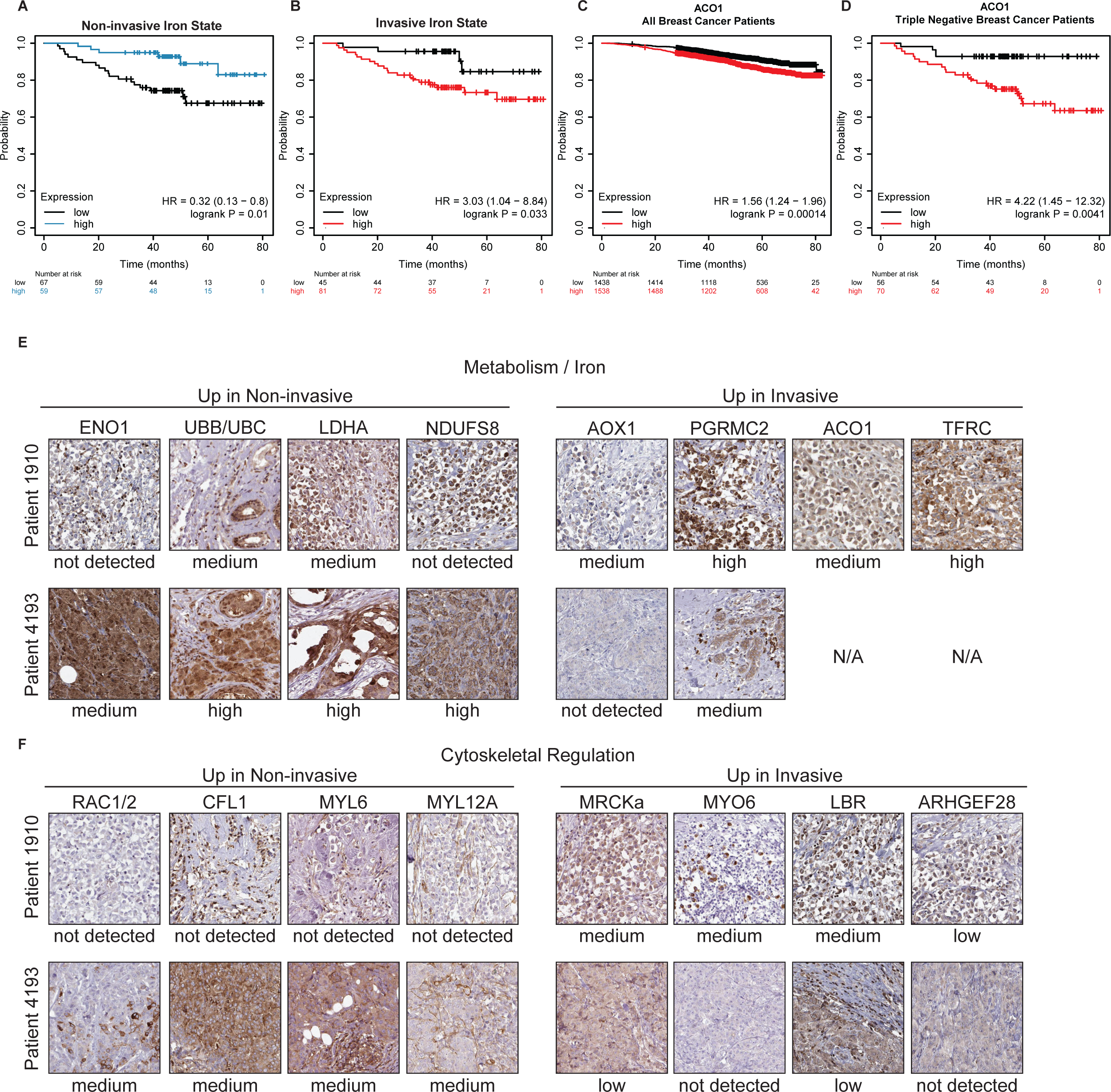
Iron-associated transcriptional states induced by HD Col1 are evident in patient tumors. **(A-B)** Kaplan Meier plots showing TNBC patient five-year overall survival stratified by high or low mean expression of the top six iron-related genes enriched in the (A) non-invasive or (B) invasive populations. **(C-D)** Kaplan Meier five-year overall survival plots for (C) all breast cancer patients or (D) TNBC patients stratified by *Aco1*. **(E-F)** Breast cancer tumor sections of Patient 1910 and Patient 4193 from the HPA show gene expression patterns matching the (E) divergent metabolism and (F) cytoskeletal regulation programs identified from the collective non-invasive and invasive phenotypes. Protein expression from Patient 1910 is consistent with the invasive phenotype, while Patient 4193 is consistent with the non-invasive phenotype. Staining and pathological analysis of HPA tissue sections is conducted by board-certified pathologists and provided on the HPA website.

Pathologist-annotated histopathology of breast cancer patients obtained from the Human Protein Atlas also revealed protein expression profiles consistent with the cytoskeletal, iron, and metabolism gene modules of invasive and non-invasive cells (Fig. 3 E-F). Tumor sections from Patient 1910 revealed a protein expression pattern consistent with the invasive phenotype, while sections from Patient 4193 are strikingly similar to the non-invasive phenotype. The architectures of these two tumors also appear to mimic the *in vitro* phenotypes, which further supports the functional relevance of the distinct cytoskeletal programs. The linear chains of cells in the tumor of patient 1910 are reminiscent of the networks formed by collectively invasive cells *in vitro*, while the round cell clusters in the tumor of patient 4193 are reminiscent of the spheroids formed by non-invasive cells *in vitro*. These results suggest that the transcriptional and phenotypic heterogeneity that emerges within populations of TNBC cells after exposure to HD Col1 *in vitro* mimics clinically relevant tumor cell states in humans.

### Dense collagen I induces Heme Oxygenase 1 activity that modulates the iron response pathway

These findings begged the question of how HD Col1 initiates the divergence of iron regulatory states in TNBC cells. Since the ECM can serve as a reservoir for some bioactive molecules, we wondered whether collagen could chelate iron in a density dependent manner, causing some cells to sense low iron availability in HD Col1 that could lead to IRP1-mediated gene expression. To determine the effect of Col1 density on the availability of iron, we assayed the iron levels in media following incubation with LD Col1 (2.5mg/ml) or HD Col1 (6.0mg/ml). No significant differences in iron content were observed between Col1 concentrations after 6hr, 12hr, or 24hr (Fig. S3).

Previous studies have shown that high density HD Col1 is unique in its ability to induce the collective invasion phenotype compared to low density (LD) Col1 environments (20–22) and that cells embedded in HD Col1 initially experience oxidative stress mimicking a low adhesion state (27). So we next hypothesized that comparing transcriptional response differences between cells in these two environments at an early time point, before phenotypic divergence was notable in HD Col1, may implicate driving factors. Bulk RNA-sequencing of MDAs in LD Col1 and HD Col1 at 24hrs was analyzed specifically in terms of iron-related gene expression patterns. *Hmox1,* which encodes for heme oxygenase 1 (HO-1), was among the most highly upregulated genes in HD compared to LD (Fig. 4 A).

**Figure 4:**
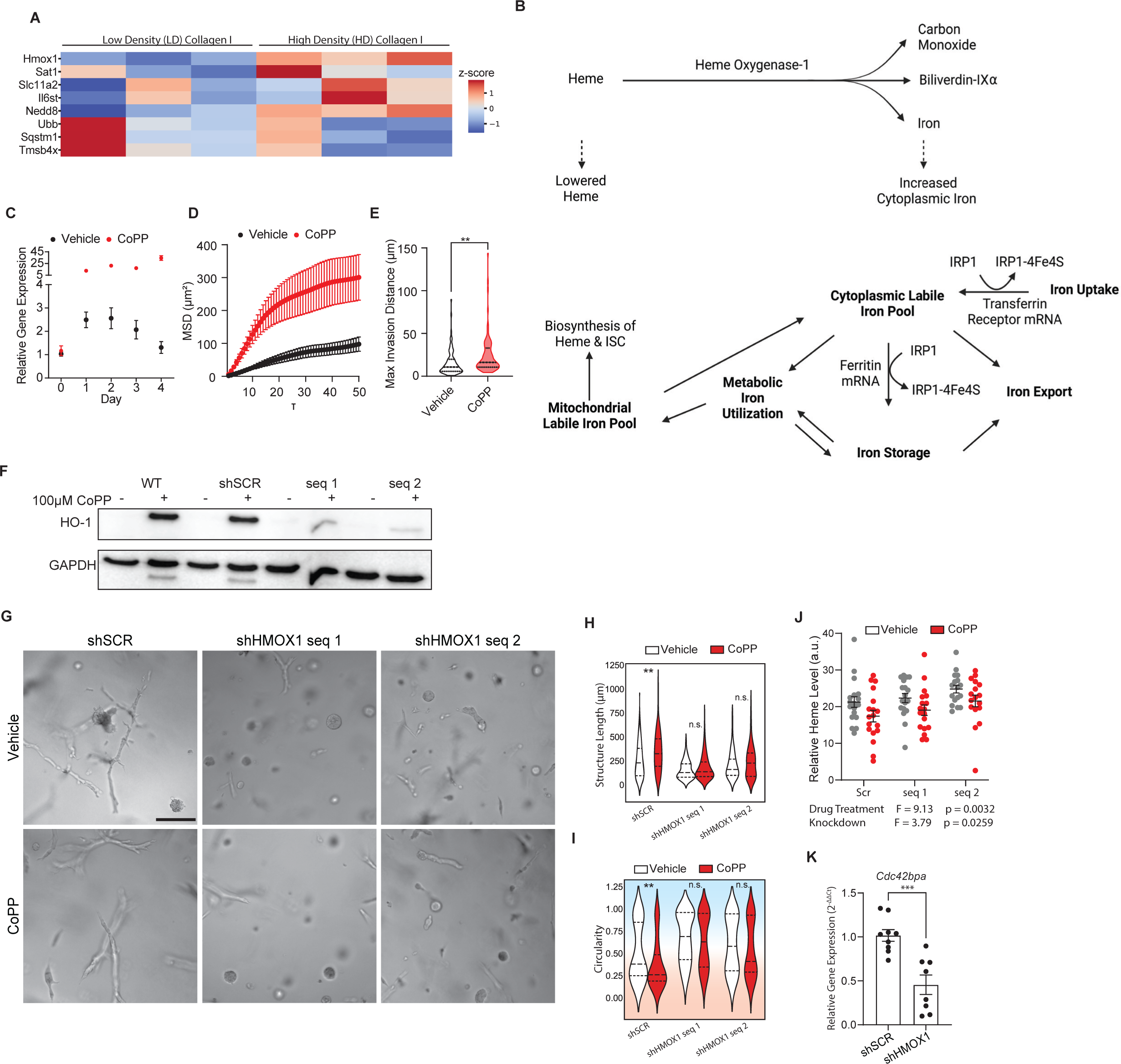
Dense collagen I initiates a low-iron response mediated by upregulation of *Hmox1*. **(A)** Bulk RNA-sequencing of MDA-MB-231 cells embedded in 2.5mg/ml (LD) or 6mg/ml (HD) collagen I matrices for 24hr reveals differential expression of iron-related genes. Cells in the HD matrices are enriched for genes related to iron import and usage, while those in LD matrices are enriched for genes related to iron sequestration or storage. Genes are ordered by fold change. Color gradient corresponds to z-score of the transcripts per million (TPM) for each gene in the two conditions. Statistical significance between the LD and HD conditions was determined using paired student’s t-tests and only genes with p < 0.05 are shown. N = 3 biological replicates per condition. **(B)** Schematic of heme oxygenase-1 enzymatic activity and the resulting effects on intracellular iron and heme levels. Adapted from (109). **(C)** RT-qPCR analysis of *Hmox1* gene expression in HD Col1 matrices shows a peak between 24-72hr for MDA-MB-231 cells. Treatment with 10μM cobalt protoporphyrin (CoPP) induces *Hmox1* expression approximately 25x higher than levels measured 1hr after embedding. *Hmox1* gene expression normalized to *Actb* transcript levels. **(D-E)** Cells treated with CoPP compared to vehicle controls are significantly more motile (D) and invasive (E). N = 3 biological replicates with n = 30 trajectories per replicate. **(F)** Knockdown of *Hmox1* confirmed by western blot. **(G-I)** Knockdown of *Hmox1* results in less collective cell invasion, while CoPP treatment enhances collective invasion only for the scramble control population. N ≥ 3 biological replicates for each condition scale bar = 50μm for trajectories and 200μm for images. Statistical significance was determined by one-way ANOVA followed by Dunnett post-test for (D) and (H), and two-way ANOVA followed by Sidák post-test for (J) and (L). * p < 0.05, ** p < 0.01, *** p < 0.001. **(J)** Heme levels in MDAs stably expressing shSCR or shHMOX1 treated with vehicle or CoPP. One-way ANOVA shows a significant effect of the knockdown and drug treatment. N ≥ 3 biological replicates for each condition. **(K)** RT-qPCR shows a down-regulation of the MRCK-encoding gene *Cdc42bpa* in the shHMOX1 cells compared to shSCR control after seven days in HD Col1. N ≥ 3 biological replicates for each condition. Statistical significance determined by student’s t-test. *** p < 0.001

HO-1 is a general stress response protein that serves as the rate-limiting enzyme in the conversion of heme into free iron, biliverdin, and CO. By cleaving heme, HO-1 contributes to the cytoplasmic labile iron pool (cLIP), and several studies have shown that iron released into the cytoplasm by HO-1 activity triggers iron sequestration gene expression through IRP-mediated pathways (50–52), mirroring the response of the non-invasive subpopulation of TNBC cells in HD Col1 (Fig. 4 B). This led us to hypothesize that the extent to which HD Col1 induced HO-1 expression in individual cells may dictate their cytoplasmic iron levels and thereby their phenotype, with high levels of HO-1 promoting high cytoplasmic iron and a non-invasive phenotype. HO-1 upregulation by cells in HD Col1 was validated by RT-qPCR. *Hmox1* expression peaked between 24-72 hrs post-embedding and fell back to initial levels by day 4 (Fig. 4 C). To test this, the heme analog cobalt protoporphyrin (CoPP) was added to MDA cells in HD Col1 to further upregulate and sustain Hmox1 expression (Fig. 4 C) and live-cell tracking was conducted to monitor migration. Counterintuitively, CoPP treatment was associated with a significant increase in the mean squared displacement (MSD) and maximum invasion distance of cells (Fig. 4 D, E). To further investigate, stable knockdowns of HO-1 were generated via lentivirally transduced shRNAs (Fig. 4 F). Embedding HO-1 knockdown (KD) cells in HD Col1 resulted in significantly fewer collectively invasive structures than the scramble controls (Fig. 4 G-I). As a control, we noted that CoPP treatment did not increase invasion in HO-1 KDs. To further confirm that these perturbations were impacting the activity of HO-1 in the intended ways, cellular heme levels were measured. As anticipated, inducing *Hmox1* with CoPP in control cells lowered heme levels (F = 9.13, p = 0.0032) and *Hmox1* KD increased heme levels compared to shSCR control (F = 3.79, p = 0.0259) (Fig. 6 J). Taken together, these results paradoxically suggest that the iron-generating activity of HO-1 drives a low cLIP state and invasive phenotype.

**Figure 5:**
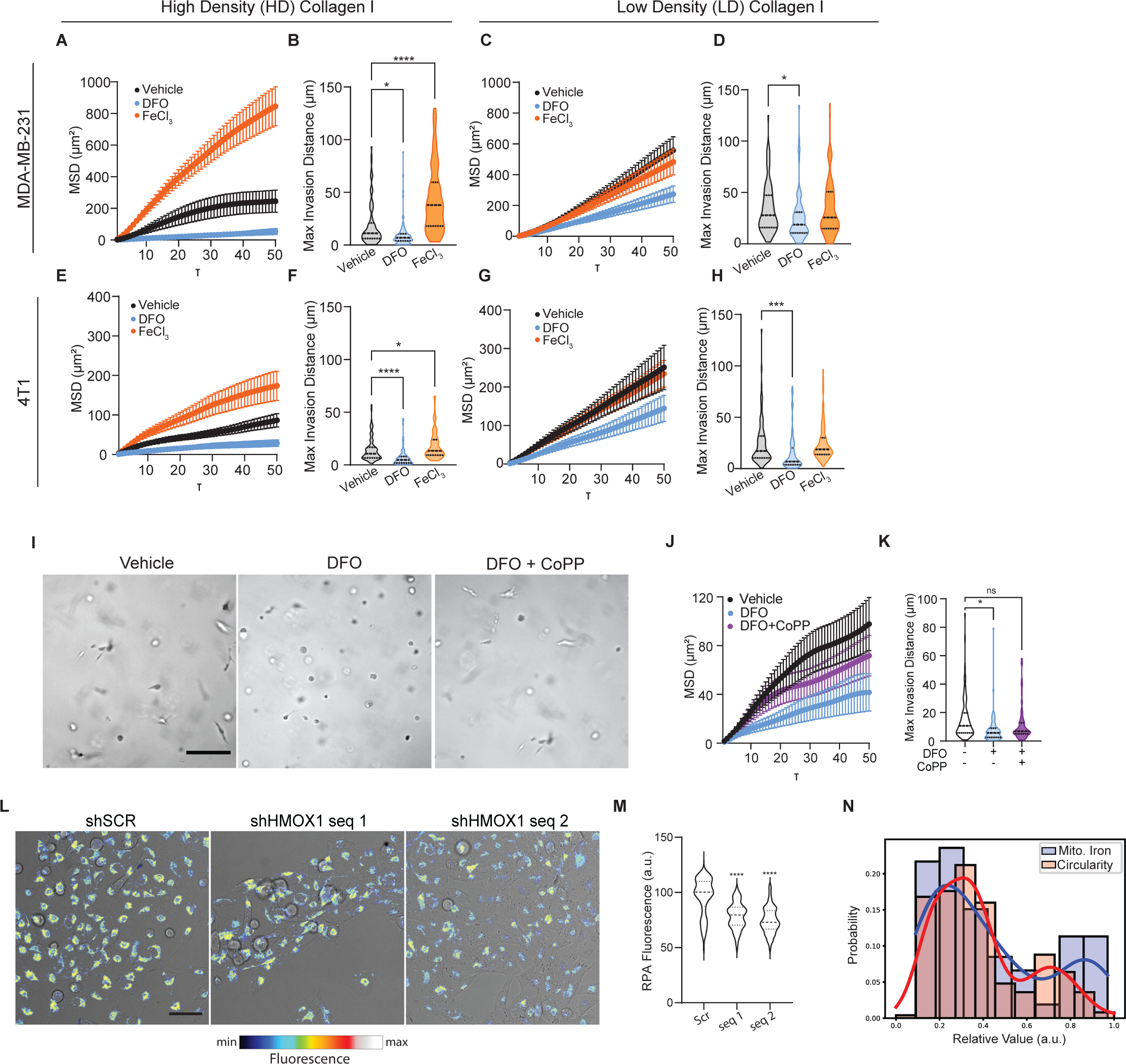
Increased iron demand promotes invasive cell migration in a collagen density-dependent manner. MSD and max invasion distances from cell tracking of MDAs in (A-B) HD Col1 and (C-D) LD Col1. **(E-H)** MSD and max invasion distances from cell tracking of 4T1s in (E-F) HD Col1 and (G-H) LD Col1. Cells were treated with vehicle (0.1% dH_2_0), 10μM DFO, or 200μM FeCl_3_. N = 3 biological replicates with at least n = 30 cells per replicate. Scale bar = 50μm. Statistical significance was determined using one-way ANOVA followed by a Dunnett post-test. * p < 0.05, ** p < 0.01, *** p < 0.001, **** p < 0.0001 **(I)** Brightfield images of MDAs after 48hr culture in HD matrices treated with either DFO or DFO+CoPP show the preservation of cell elongation by CoPP. scale bar = 200μm. **(J-K)** Migration analysis shows the protective effects of CoPP treatment when cells were challenged with DFO. N = 3 biological replicates with n = 30 trajectories per replicate. (L) Representative micrographs of MDAs showing brightfield and RPA fluorescence channel overlaid of shSCR control and shHMOX1. (M) Quantification of the fluorescence within single cells shows a significant reduction in the shHMOX1 cells, indicating higher levels of mitochondrial labile iron. (N) Histogram of fluorescence levels within single cells of the shSCR cells shows a bimodal distribution of intensities consistent with the proportion of invasive vs. non-invasive multicellular structures when embedded in 3D Col1 matrices. Scale bar = 50um

**Figure 6:**
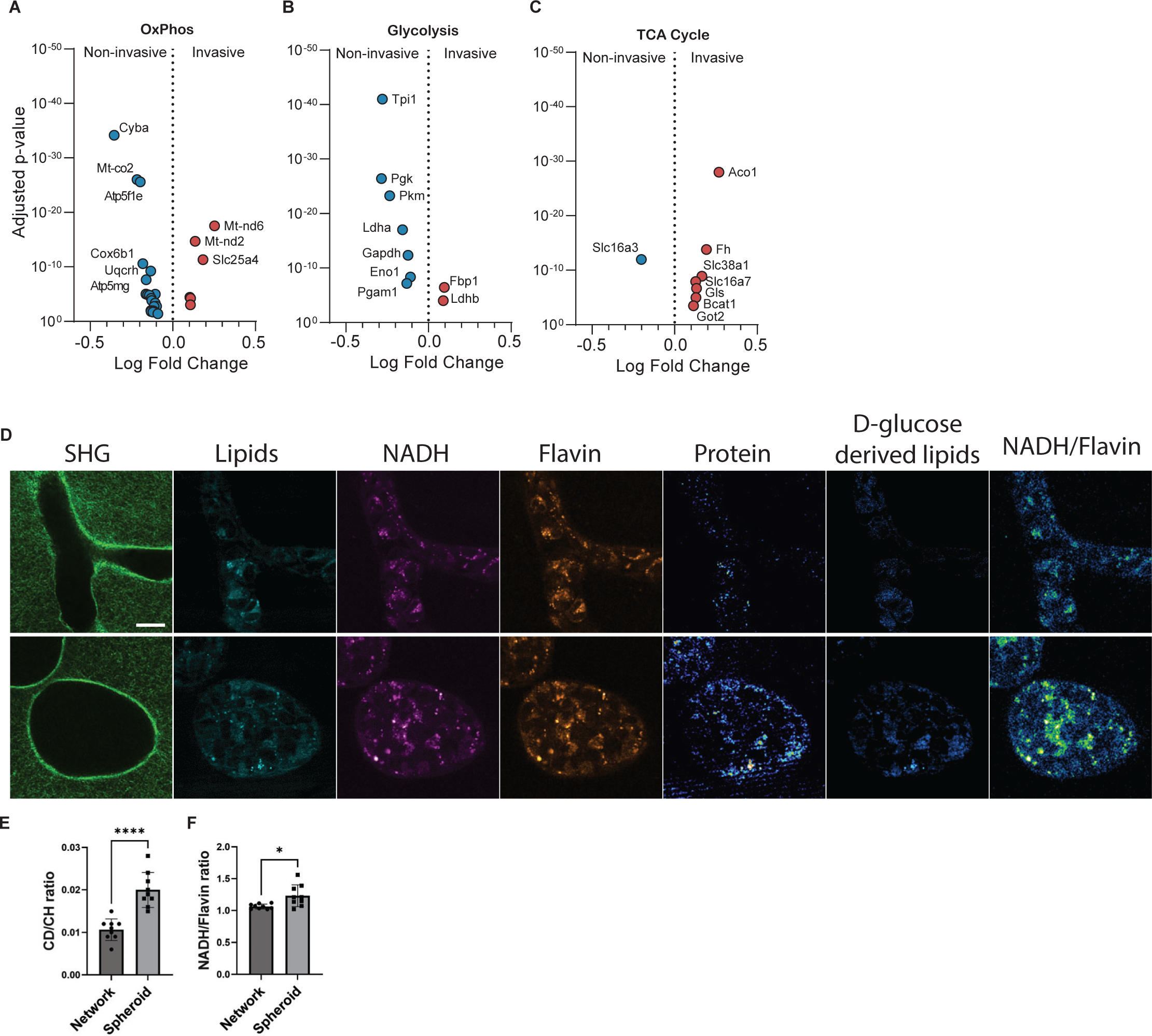
Non-invasive collective migration is more glycolytic than the more OxPhos-dependent invasive collective migration phenotype. **(A-C)** Differentially expressed genes related to (A) OXPHOS, (B) glycolysis, and (C) TCA cycle. **(D)** Representative micrographs from stimulated raman spectroscopy of MDA collective migration in HD Col1. **(E)** Quantification of the ratio of CD:CH shows an increase in the non-invasive structures, indicating higher glycolytic flux. **(F)** The ratio of NADH/Flavin is significantly lower in the invasive structures, indicating more flux through the TCA and OxPhos pathways. Statistical significance determined by student’s t-test. * p < 0.001; **** p < 0.0001

To further validate the relationship between HO-1 activity and the IRP-mediated cLIP sensing pathway, RT-qPCR analysis of the IRP1 regulated MRCK-encoding gene *Cdc42bpa* was conducted on shSCR versus shHMOX1 cells after seven days in HD Col1. HO-1 KD lowered expression of *Cdc42bpa* compared to shSCR cells. Since *Cdc42bpa* expression increases in response to low cLIP conditions via IRP regulation, and HO-1 expression increases *Cdc42bpa* expression, this further suggests that HO-1 can promote a low cLIP response state that determines migration outcomes (Fig. 4 K).

### HO-1 drives a low cytoplasmic iron response by inducing high iron utilization and OXPHOS

The iron level of the cLIP is balanced by uptake, export, storage, and metabolic utilization. We posited that for cLIP levels to be low despite the release of iron from heme by HO-1, invasive cells may be utilizing iron at a rate that outpaces heme cleavage. This would suggest that the invasive state represents a high iron flux and consumption state rather than simply a static low cytoplasmic iron state. To begin to test this, we asked how exogenous iron levels impact invasion. The MSD and max invasion distance of MDAs in HD Col1 supplemented with extracellular iron in the form of FeCl_3_ were significantly higher. Likewise, iron chelation dramatically reduced their migration (Fig. 5 A, B, S4). However, MDAs in LD Col1 were unaffected by increasing iron levels and iron chelation decreased motility to a lesser extent than in HD Col1 (Fig. 5 C, D, S4). Similar patterns of response were observed for 4T1 cells (Fig. 5 E-H, S4). This confirmed that TNBC cells within dense collagen I matrices become more dependent on iron and utilize iron more to facilitate migration. However, the effects of DFO were somewhat surprising. The anti-cancer effects of DFO are not mediated by prevention of iron uptake by transferrin but rather by an intracellular pool of iron that is necessary for DNA synthesis (53). Therefore, we hypothesized that further *Hmox1* induction, beyond that triggered by HD Col1, could counteract the anti-migratory effects of DFO chelation. To test this, we chelated iron using DFO and simultaneously treated cells with CoPP. The addition of CoPP treatment did counteract the effect of DFO on migration and invasion (Fig. 5 E-G, S4). CoPP intervention not only rescued cell migration, but also restored the population of cells with elongated morphologies (Fig. 5 E). This confirmed that HO-1 induction provides an important endogenous source of iron that TNBC cells can use to fuel an invasive state with high iron utilization requirements.

Iron utilization primarily takes place in the mitochondria where it is used for heme and ISC synthesis by interdependent and coordinated processes and incorporated into mitochondrial proteins that support essential functions like respiration (54, 55). Accordingly, the invasive cell state is associated with the upregulation of several ISC-containing genes (*Abce1*, *Hspa9*, *Brip1*, *Aox1*, *Rrm2*, *Adi1*, *Cdkal1*, *Ppat*, *Dpyd*, *Glrx5*, *Dna2*, *Prim2*) that are necessary for processes like DNA repair, RNA degradation, protein folding, and cellular metabolism. Additionally, *Pgrmc2* is a crucial heme transporter between the cytosol and mitochondria (Fig. 2). To assess whether the HO-1 mediated increase in iron utilization is linked to mitochondrial iron usage, we measured mitochondrial chelatable iron levels upon perturbation of HO-1. Increases in mitochondrial biosynthetic activity have been shown to have the effect of lowering chelatable mitochondrial iron levels (56). CoPP was used to robustly induce *Hmox1* expression in MDAs in 2D culture conditions to mimic the effect of HD Col1 on *Hmox1* expression. *Hmox1* induced shSCR cells had lower levels of chelatable iron in their mitochondria (higher RPA fluorescence, Fig. 5 L, M) than shHMOX1 cells (lower RPA fluorescence, Fig. 5 L, M), which is consistent with increased mitochondrial iron utilization in response to *Hmox1* induction. Interestingly, the chelatable mitochondrial iron levels measured in scramble control cells reveal two sub-populations: ∼70.4% have low chelatable mitochondrial iron (high RPA fluorescence) and 29.6% show high chelatable mitochondrial iron (low RPA fluorescence) (Fig. 5 N). These proportions are similar to the distribution of invasive and non-invasive cells (Fig. 5 N), providing further evidence that the invasive phenotype represents a high mitochondrial iron utilization phenotype.

Increased heme synthesis by mitochondria has also been shown to increase energy production through oxidative phosphorylation (OXPHOS) (57–59), as heme is an essential component of OXPHOS complexes II–IV (60) and is required for ATP production by the mitochondrial ETC and OXPHOS (61). This led us to re-analyze the phenotypic gene expression profiles specifically for DE of glycolysis, OXPHOS, and the TCA cycle components. Multiple genes involved in OXPHOS were differentially expressed between the two phenotypes (Fig. 6 A, Fig. S5, but glycolysis-related genes were preferentially upregulated in the non-invasive cell population (Fig. 6 B, Fig. S5). Strikingly, nearly every enzyme that promotes the catabolism of glucose to lactate was enriched in the non-invasive phenotype. The only glycolysis-related genes upregulated in the invasive phenotype were *Fbp1*, which catalyzes a reverse step of glycolysis, and *Ldhb*, which preferentially promotes the conversion of lactate back to pyruvate (62). In contrast, the invasive cells upregulated genes involved in the TCA cycle (Fig. 2 E, Fig. S5). These genes suggest a program of anaplerotic fueling of the TCA cycle with glutamate via glutamine import and conversion (*Slc38a1*, *Gls*, *Got2*). These data suggest that the non-invasive phenotype is more dependent on glycolysis while the invasive phenotype is more dependent on OXPHOS. To test this, we used stimulated Raman-scattering (SRS) to image glucose metabolism in invasive and non-invasive cells. SRS spectral tracing of deuterated glucose (D-glucose) revealed higher levels of glycolysis-derived biosynthesis in non-invasive structures (Fig. 6 D, E), and NADH/FAD redox ratio measurements were higher on average in non-invasive structures than invasive structures (Fig. 6 D, F). These data confirm that invasive cells rely more on OXPHOS while non-invasive cells rely more on glycolysis.

Given these fundamental metabolic dependencies, we hypothesized that each cell phenotype would be differentially susceptible to perturbations of glycolysis and OXPHOS. We also noted that the glycolysis enzyme ENO1, which was transcriptionally enriched in the non-invasive phenotype and has been a target of interest for cancer drug development, has recently been shown to have an iron-regulatory role via modulating the degradation of ACO1 (63). Recall, ACO1 was transcriptionally upregulated in the OXPHOS-dependent invasive phenotype and was necessary for invasion (Fig. 2A, C-F). Motivated by this antagonistic relationship, we challenged both MDA and 4T1 cells with inhibitors targeting ENO1, to diminish glycolysis, or with inhibitors of Complex I, which mediates OXPHOS (Fig. 4 F). Complex I inhibitor rotenone stunted collective invasion, decreasing cell structure lengths (Fig. 7 B, Fig. S6 A-C) and increasing their circularity (Fig. 7 C, Fig. S6 D-E). Conversely, ENO1 inhibitor AP-III-a4 slightly increased the proportion of long structures and decreased the number of circular structures (Fig. 7 B, C, Fig. S6 A-E). Both Complex I and ENO1 inhibition decreased cell viability, though to different extents (Fig. 7 D). At 1µM rotenone (IC50 ∼1.7µM (64, 65)), cell viability was 37.89%, while ENO1 inhibition at 1µM (IC50∼0.6µM (66)) led to approximately 79.82% viability. Together, these data suggest that inhibition of glycolysis predominantly eliminated the non-invasive population, while inhibition of OXPHOS primarily eliminated the invasive population.

**Figure 7:**
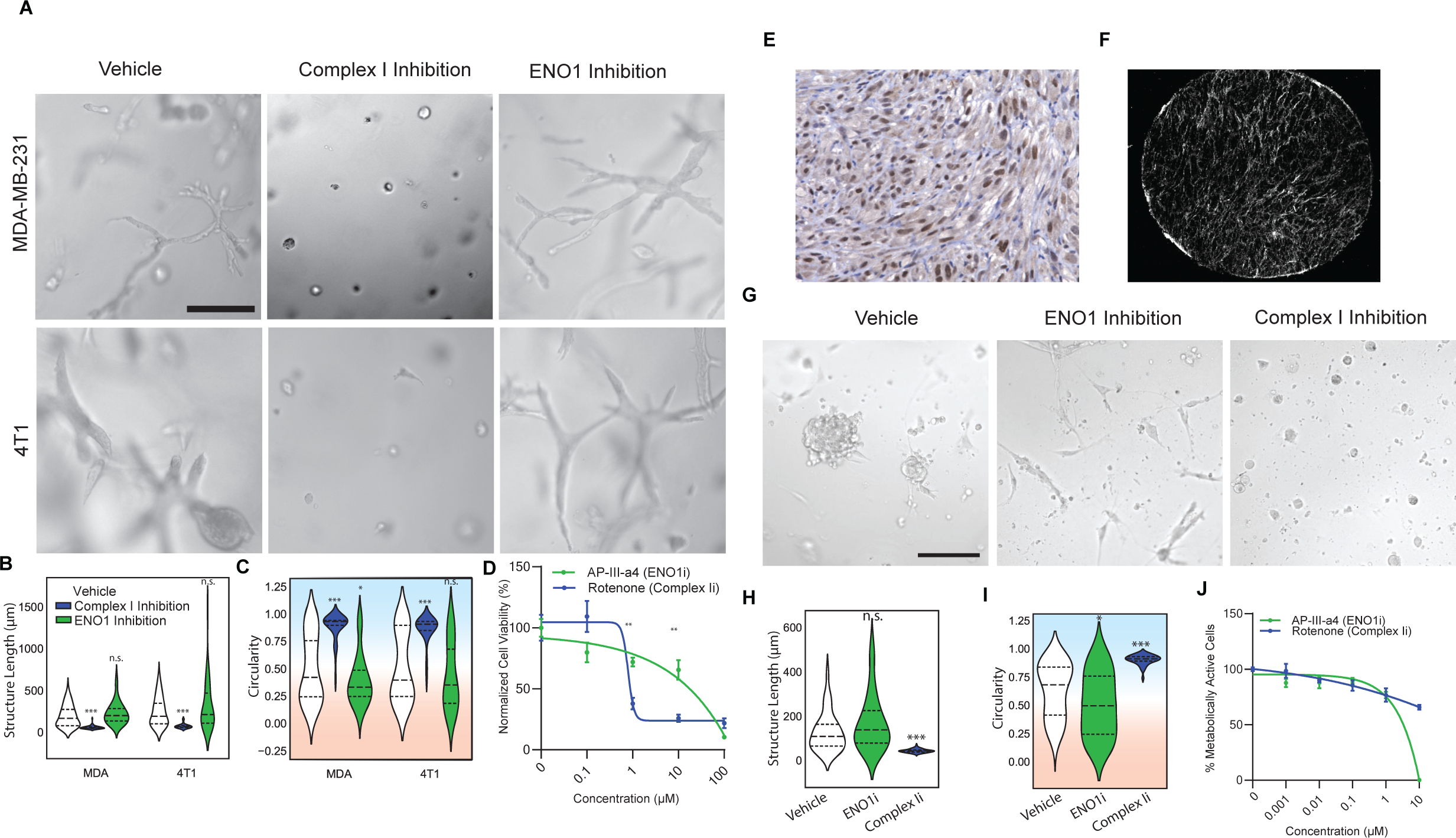
Collectively invasive cells are more sensitive to OxPhos inhibition than non-invasive cells. **(A)** Representative micrographs of MDAs in HD Col1 treated with 0.1% DMSO vehicle, 1μM complex I inhibitor rotenone, or 1μM glycolytic enzyme ENO1 inhibitor AP-III-a4. Inhibition of complex I abrogated the collective invasion phenotype, while inhibition of ENO1 decreased the proportion of the non-invasive phenotype in both MDAs and 4T1s. **(B-C)** Quantification shows the maximum length of structures (Feret Diameter) (B) and circularity (C). N = 3 replicates for each condition. Individual structures for MDAs: n = 70 vehicle, n = 71 ENO1i, n = 64 Complex Ii. 4T1s: n = 56 vehicle, n = 42 ENO1i, n = 36 Complex Ii. **(D)** MTS assay results showing relative cell viability in response to drug treatments. **(E)** Representative H&E histology section of the PA-1413 patient-derived organoid model showing a strand of collectively invasive cells. Counterstaining for TWIST1 is shown in brown. **(F)** Representative second-harmonic generation (SHG) image of the dense fibrous collagen architecture of the PDO. **(G)** Representative micrographs of patient-derived organoid models treated with 0.1% DMSO vehicle, 1μM complex I inhibitor rotenone, 1μM glycolytic enzyme ENO1 inhibitor AP-III-a4, or MRCK-inhibitor BDP-9066. Scale bar = 100μm **(H-I)** Quantification shows the maximum length of structures (Feret Diameter) (K) and circularity (L). **(J)** MTS assay results showing relative cell viability in response to drug treatments. Scale bars = 200μm. N = 3 biological replicates per treatment group. Statistical significance was determined using one-way ANOVA followed by a Dunnett post-test comparing each treatment group to the vehicle group. * p < 0.05, ** p < 0.01, *** p < 0.001, **** p < 0.0001.

To validate the culmination of our findings in a system that is considered to be more clinically relevant, we identified a patient-derived organoid (PDO) model of TNBC (PA-14-13) whose microenvironment is rich in Col1 and whose phenotype resembles the collective chains of cells formed in our *in vitro* system and in patient tumors expressing the invasive cell state (Fig. 7 E, F). Applying ENO1 inhibitor to PA-14-13 PDOs resulted in the prevalence of organoid cells with an elongated phenotype compared to vehicle conditions (Fig. 7 G-I). Complex I inhibition resulted in the prevalence of organoid cells with a circular phenotype and largely eliminated the elongated phenotype (Fig. 4 G-I). These effects were observed at 1uM concentrations where viability at 76.85% for ENO1 inhibition and 76.84% for Complex I inhibition (Fig. 7 J). These findings echo the results of our HD Col1 model system and further support the concept that phenotypic heterogeneity in collagenous TNBC tumors is driven by divergent metabolic dependencies.

## Discussion

Despite a wealth of data linking breast cancer to iron and a significant interest in iron chelators and iron overload as possible anticancer strategies (67) (68, 69), the mechanisms driving dysregulated iron metabolism in breast cancer remain largely uncharacterized or suffer from conflicting reports in the literature. Our study contributes several findings to advance the current state of knowledge in this area. First, it reveals that TNBC cell iron dependency is regulated by ECM conditions. Dense collagen type I shifts the majority of TNBC cells within a population into a high iron utilization state that drives invasive behaviors and sensitizes TNBC cells to iron chelation. Exogenous iron promoted the invasion of TNBC cells in HD Col1 significantly more than in LD Col1, and chelation of iron with DFO inhibited the invasion of TNBC cells in HD Col1 to a greater extent than in LD Col1. Some other reports of the effects of DFO on MDAs are consistent with our findings (70), while some previous reports contradict these results (71, 72). An important distinguishing factor is that the effects observed in HD Col1 are within the context of ECM-induced HO-1 activity.

How exactly dense Col1 stimulates HO-1 is unclear. HO-1 induction usually indicates the mobilization of the cellular antioxidant response to oxidative stress, and our prior studies have shown that at early time points after cancer cells are embedded in dense Col1, they show signs of oxidative stress linked to low adhesion (27). Alternatively, others have shown that reactive oxygen species (ROS) can be generated by the rupture of Col1 under tensile loads (73). The ROS generated by Col1 rupture can be converted to hydrogen peroxide or quinones; the former is a potent cell signaling molecule and the latter may induce collagen to form complexes with iron (74). Both species could explain the HO-1 response.

HO-1 has been shown to have a dual role as antitumor and protumor in several cancers including breast(75–82). Our study reveals one way in which HO-1 induction can result in the emergence of two distinct iron regulatory states and metabolic states that are associated with opposite patient outcomes. These states mimic to some extent the canonical IRP1-mediated gene expression responses to high and low cLIP conditions. The state resembling the response to high cLIP levels mirrors the iron storage transcriptional response of several normal cell types to HO-1 (48), relies more on glycolysis, and predicts better prognosis. However, the state that resembles the transcriptional response to low cLIP levels and predicts poor prognosis occurs in conjunction with a low mLIP state and reliance on OXPHOS, which is characteristic of increased mitochondrial biosynthesis activity (56) (57–59). This state may be best described as a high iron utilization state that maintains a low cLIP response. Of note, this gene set overlaps to some extent with previously published iron-regulatory gene sets that predict poor prognosis in breast cancer patient cohorts (83). Since cytoplasmic iron homeostasis and mitochondrial iron homeostasis are interdependent (84), HO-1 activity may achieve this state by simultaneously contributing iron to the cLIP while lowering heme levels, which could initiate heme synthesis (85) and lower the mLIP, increasing iron flux to the mitochondria. HO-1 KD increased cellular heme levels and abrogated collective invasion in HD Col1, which aligns with other studies showing that increasing heme levels decreases TNBC cell migration (86). Likewise, several recent reports suggest that elevated heme synthesis underlies enhanced OXPHOS and tumorigenicity in TNBC, ovarian cancer, and non-small cell lung cancer, and pancreatic cancer (59, 87, 88). Our functional studies suggest that ACO1/IRP1, MRCK, and OXPHOS could be targets of interest for TNBC cells in this state, motivating future studies *in vivo*. High ACO1/IRP1 expression also strongly predicted poor survival in TNBC patients and could be investigated as a biomarker for this state. Interestingly, ACO2 has been shown to promote colon cancer progression (89). Other targets may be identified by further functional studies of the DE genes identified in this study.

While cancer cells are typically more glycolytic, and several studies have reported anti-proliferative and anti-invasive effects of glycolysis inhibition through ENO1 (90), our experiments inhibiting ENO1 in cell lines and a PDO with collagenous microenvironments show that this targeted only the fraction of the population that is non-invasive. Inhibiting OXPHOS targeted the invasive population. OXPHOS is emerging as a potential therapeutic target based on the observation that it is upregulated in certain tumors (91) and can drive metastatic behaviors (92). The relative contribution of glycolysis and OXPHOS to ATP production has been shown to be highly variable between cancer cell types (93), and recent evidence suggests that increasing Col1 density shifts cell populations away from glycolysis and towards OXPHOS (94). Our data also demonstrate this influence of Col1 density on metabolism and further show that this shift is accompanied by metabolic divergence within cell populations. This supports the emerging paradigm that a combination of microenvironmental and cell-intrinsic factors determine stable cell states and thereby context-dependent effects in tumors (13, 14, 95).

The experimental findings of this study come from 3D culture and patient-derived tumor organoids, and should therefore be considered with an understanding of the inherent limitations of these model systems. Collective cell invasion is strongly correlated with metastasis (16), though more invasive cells do not necessarily directly translate to more metastatic cells. Indeed, increasing evidence shows that the most efficient metastatic seeding occurs from heterogeneous clusters of cancer cells (19). To understand the impact of our findings on metastasis, *in vivo* studies will be required. However, the clinically relevant tumor cell states we induced by controlling the ECM *in vitro* are expected to be more difficult to reliably replicate *in vivo*. Identifying *in vivo* models that recapitulate the ECM and cellular architectures of clinically relevant TNBC cell states is an important challenge.

In summary, our findings provide new information about iron-regulatory and energy metabolism states of TNBC cells and the role of the ECM on these states through impacts on cellular iron flux. This may inform current iron-targeted treatment strategies and implicate new strategies for further investigation.

## Materials and Methods

### Cell Culture

MDA-MB-231 (RRID:CVCL_0062), 4T1 (RRID:CVCL_0125), LentiX (RRID: CVCL_4401), and LMP (96, 97) cells were cultured at 37°C and 5% CO2 in high glucose Dulbecco’s Modified Eagle Medium (Gibco Cat# 11995065) supplemented with 10% (v/v) fetal bovine serum (Corning Cat# 35010CV) and 0.1% (v/v) gentamicin (ThermoFisher Cat# 15750-060). Cells were passed when they reached 80% confluency.

### Bulk RNA Sequencing Heat Map

The RNA sequencing analyzed in this experiment was performed previously by our lab and is publicly available (21). Briefly, cells were isolated from LD and HD Col1 matrices after 24hr in culture. Total RNA was sequenced on the Illumina MiSeq platform (Illumina, San Diego), reads were aligned using Bowtie2 (RRID:SCR_016368) and eXpress, and genes were mapped to the human genome hg19UCSC. New analyses of this data were conducted for this paper to identify iron-related genes using gene lists available from MSigDB (http://www.gsea-msigdb.org/gsea/index.jsp).

### Cell Tracking

Live cell tracking was performed using a Nikon Ti Eclipse Inverted Epiflourescent microscope equipped with an incubation system to maintain 37°C, 5% CO2, and humidity. Bright field images were acquired using a 10x objective at 10 minute intervals. Cell tracking analysis was performed using Metamorph Software (RRID:SCR_002368, Molecular Devices, San Jose, CA) to generate xy-trajectories, followed by a custom MATLAB (RRID:SCR_001622, Mathworks, Natick, MA) script to calculate the migration parameters. MSDs were plotted using the first third of possible tau values. Max invasion distance was calculated as the maximum linear distance traveled from each cell’s starting position.

### Collective Migration Phenotype Quantification

Cells were cultured in the HD matrices for seven days and representative images from each individual gel were acquired on the same microscope described above. At this point the cells have formed distinct collective structures, which are outlined and shape descriptors are calculated using ImageJ (FIJI) (RRID:SCR_002285) (98). The ferret diameter of multicellular structure was used to report the invasion distance. Structures with a circularity index greater than 0.7 are considered non-invasive, while scores lower than 0.7 are considered invasive.

### Lentiviral Transduction

Lentiviral transduction of shRNA targeting *Hmox1* was completed following viral preparation using plasmids supplied by the La Jolla Institute for Immunology (Seq1: TRCN0000045250: ACAGTTGCTGTAGGGCTTTAT, Seq2: TRCN0000045251: CAACAAGGAGAGCCCAGTCTT) and gifted from Didier Trono (RRID:Addgene_12259, RRID:Addgene_12260). *Aco1* was targeted using sequences from the Mission Database (Seq1: TRCN0000056553: CCAGGAAAGAAATTCTTCAAT, Seq2: TRCN0000056554: GCAGGATTGTTAGCAAAGAAA) shRNA plasmids were cotransfected with the packaging and envelope plasmids using Lipofectamine 3000 at a ratio of 4:3:1 (#L3000001, Thermofisher) into LentiX cells. After 48hrs the media containing the lentivirus was collected and filtered through a 0.45μm filter. MDA-MB-231 cells were transduced with the lentivirus for 48hr, followed by selection with 1μg/mL puromycin ( Millipore Sigma Cat# P9620). Protein-level knockdown was validated with a western blot.

### 3D collagen gels

Collagen matrices with embedded cells were made as previously described (21). Briefly, cells were mixed with reconstitution buffer and unpolymerized rat tail collagen I ( #CB354249, Corning, Corning, NY) to reach the desired collagen concentration, followed by the addition of 1N NaOH to initiate polymerization. After 30 minutes at 37LJ, media was added onto the polymerized gels. Collagen was imaged using confocal reflection microscopy to confirm fiber architecture consistency between experiments. Media was changed every two days for the duration of the cultures. CoPP was supplied by Enzo LifeSciences (Cat# ALX-430-076-M025, Farmingdale, NY) and resuspended in DMSO (Sigma Aldrich, St. Louis, MI). DFO (Cat# D9533) and FeCl_3_ (Cat# 157740) were supplied by Sigma Aldrich (St. Louis, MI) and resuspended in dH20. Inhibitors rotenone (Cat# HY-B1756), AP-III-a4 (Cat# HY-15858), and BDP9066 (Cat# HY-111424) were purchased from MedChemExpress (Monmouth Junction, NJ). Cells were treated with the inhibitors for 24hrs, after which they were treated with normal media. Vehicle control samples were treated with 0.1% DMSO. Collagen concentrations were chosen based on previous studies that characterized both the mechanical properties and cellular behaviors of cancer cells embedded within those matrices (21, 22, 28, 29, 99).

### Western Blotting

Western blot samples were prepared as previously described. Briefly, collagen gels were washed with ice cold PBS then scooped into protein low-bind tubes and placed on ice. RIPA lysis buffer (ThermoFisher Cat# 89900) with 1x protease and phosphatase inhibitors (ThermoFisher Cat# 78446) was added at an equal volume as that of the gels. After a brief 30 second sonication on ice, loading buffer (ThermoFisher Cat# B0007) was added to the gels to reach 1x concentration. Samples were incubated on ice for an hour, with vortexing every 15 minutes. Finally, samples were incubated at 95C for 5 minutes and stored at -20C until use. Proteins were separated in 10% SDS-PAGE gels and transferred to 0.45um PVDF membranes in 20% methanol Towbin buffer. Membranes were blocked with 5% BSA and incubated with primary antibodies for HMOX1 (Abcam Cat# ab52947, RRID:AB_880536), ACO1 (Santa Cruz Biotechnology Cat# sc-166022, RRID:AB_2273699), or GAPDH (Cell Signaling Technologies Cat# 2118S) overnight at 4LJ. Washing was done 3x with TBST before and after secondary antibodies (Cell Signaling Technology Cat# 7074, RRID:AB_2099233) were added. ECL was added before visualization in a gel reader.

### Single cell RNA Sequencing

The phenotypically-supervised single-cell RNA sequencing data used in this study was generated previously by our lab and is publicly available (28). Briefly, cells expressing the photo-convertible fluorescent protein Dendra-2 Lifeact were imaged after 7 days and the phenotype of interest was converted from GFP to RFP. Cells were then isolated and sorted by FACS prior to sequencing. scRNA-seq was performed using the Illumina NovaSeq 6000 and analyzed using the cellRanger pipeline, aligning to reference human genome GRCh38.

### Pathway enrichment analysis

Differentially expressed genes were determined using the non-parametric Wilcoxon rank sum test with a Bonferroni adjusted p-value < 0.05. Pathway enrichment analysis on the 1,706 DE genes was performed using the Reactome 2022 database on the Enrichr platform (100–102). The “combined score” enrichment value is calculated by taking the log of the p-value from the Fisher exact test and multiplying that by the z-score of the deviation from the expected rank.

### RT-qPCR

MDA-MB-231 cells were embedded in 200uL HD Col1, then samples were treated with equal volume of Trizol (#15596026) at the specified timepoints. RNA was extracted using the Roche High Pure RNA Isolation Kit (#11828665001), and cDNA was created using SuperScript III reverse transcriptase (#18080044, Invitrogen). Taqman probes for *hmox1* (Hs01110250_m1) and *actb* (Hs01060665_g1) were used for qPCR in a QuantStudio3 thermocycler (ThermoFisher). Relative gene expression was calculated from 2^-□□Ct^ relative to vehicle control on Day 0.

### Kaplan-Meier Plots

Kaplan-Meier plots were generated using the online Kaplan-Meier Plotter tool (kmplot.com) (49). This tool manually curates patient survival data from TCGA, EGA, and GEO and combines gene expression with clinical data. All analyses used RNA-seq data of breast cancer patient overall survival. The mean expression of the top six iron-related genes in the non-invasive and invasive phenotypes for triple negative breast cancer patients was used for the plots in Figure 1. Analyses for *Aco1* expression for overall survival were run on RNA-seq data on two groups of patient samples split by quantile expressions. Hazard ratio is shown with 95% confidence interval, along with the logrank P value.

### Cancer Patient Histological Data

Histology of breast cancer patients was obtained by searching the Human Protein Atlas (HPA) online database for proteins encoded by differentially expressed genes identified using PhenoSeq (https://www.proteinatlas.org/humanproteome/pathology) (103). Patient 1910 and Patient 4193 were selected for analysis because they were stained for many of the same proteins, which also were relevant to our results. Antibodies that were not specific to different isoforms are labeled for both possible proteins, which was the case for UBB/UBC and RAC1/RAC2. Patient 4193 did not have sections stained for TFRC or ACO1. Staining and pathological analysis of HPA tissue sections is conducted by board-certified pathologists and provided on the HPA website. The grades assigned in Figure 2 J-K are the direct HPA annotations.

### Patient-derived Organoid Culture and Imaging

PA-14-13 Patient-Derived Organoids were cultured in Advanced DMEM/F12 (Gibco), supplemented with 5% fetal bovine serum (FBS, Corning), 10ng/mL hEGF, 10mM HEPES, 1µg/mL hydrocortisone (Sigma-Aldrich), and 0.1% gentamicin (Thermo Fisher Scientific). All patient-derived organoids were maintained at 37°C and 5% CO_2_ in a humidified environment.

Patient-derived xenograft models using de-identified tumor samples were previously established by Helen Piwnica-Worms (MD Anderson) (104). Generation of patient-derived organoids were adapted from the following protocols (104, 105). Briefly, single cells of a PA-14-13 tumor were engrafted into each of the left and right flanks of mammary fat pads of NOD/SCID mice and when tumors reached approximately 1cm^3^, they were harvested, pooled and dissociated into smaller clusters of cells, without digesting all the way into single cells. This was intended to retain components of the native tumor architecture. First, excised tumor samples were washed in HBSS and finely minced before digesting in 50 mL of digestion media for 1 hour, which composed of Advanced DMEM/F12 (Gibco) supplemented with 2% Bovine Serum Albumin, 5µg/mL insulin, 0.2% trypsin, 0.1% gentamicin (Thermo Fisher Scientific), and 2mg/mL collagenase (Roche). Following digestion and three subsequent PBS washes, tumors were centrifuged for 10 minutes at 1500 rpm to remove fatty components and aqueous layers. Digested PDX tumor cells were treated with 1 mg/mL DNase I (Stemcell Technologies) for 1-2 minutes until sample lost viscosity. Cells were then resuspended in 50 mL media and went through five quick (<5s) 1500 rpm centrifugation steps to remove all mouse fibroblasts. Finally, PDX tumor cells were resuspended in 2% Matrigel (Corning) in media with 0.1% DMSO, Rotenone, AP-III-a4, or BDP-9066 at the indicated concentrations and seeded on top of a 60 µL 6 mg/mL Collagen I gel cultured in a 96-well plate in a sparse seeding density (20-30% confluency). Patient organoids were treated with 100µL vehicle or test compounds for 72 hours. An MTS assay was conducted to measure metabolically active cells, as described (See MTS Assay in Methods).

Immunofluorescence staining was conducted to validate depletion of mouse cells and assess tumor content in PDO cultures, using the anti-human EpCAM antibody (HEA-125, Miltenyibiotec). Briefly, wild-type PA-14-13 PDO cultures (72 hours) were washed with PBS and fixed for 30 minutes at room temperature in 4% paraformaldehyde (PFA) solution. MDA-MB-231 (human triple negative breast cancer) and LMP (murine pancreatic ductal adenocarcinoma) cells were seeded on top of a 60 µL 6 mg/mL Collagen I gel at seeding density of 50000, fixed the following day, and used as positive and negative controls for the anti-human EpCAM antibody, respectively. The fixed samples were washed again with PBS and PBS supplemented with 100mM glycine. Blocking was performed overnight at 4LJ in a solution of 1% (w/v) BSA, 2.5% FBS (v/v), 2.5% (v/v) goat serum, and 0.1% (v/v) Triton-X. Anti-EpCAM-FITC primary antibody was added at a 1:50 dilution and incubated at 4°C for 48hr, then washed out with PBS overnight. Imaging was performed on a Nikon Ti Eclipse Inverted Epiflourescent microscope.

### Rac1 Immunofluorescence Imaging

WT MDA-MB-231 cells were cultured in HD Col1 as described above in the “3D Collagen Gels” section. After seven days in culture, the gels were washed three times with ice cold PBS and fixed for 30 minutes at room temperature in 4% paraformaldehyde (PFA) solution. The fixed samples were subsequently washed again with PBS and PBS supplemented with 100mM glycine. Blocking was performed overnight at 4C in a solution of 1% (w/v) BSA, 2.5% FBS (v/v), 2.5% (v/v) goat serum, and 0.1% (v/v) Triton-X. Rac1 primary antibody was added at a 1:50 dilution and incubated at 4C for 48hr, then washed out with PBS overnight. Secondary antibody diluted 1:1000 was added overnight. Subsequent PBS washes were performed, where DAPI and phalloidin were added for 15 minutes at RT for nucleus and actin references. Imaging was performed on a Leica SP8 confocal microscope at 40x. Z-stacks were taken through 5-10 invasive and non-invasive multicellular structures for each sample.

### MTS Assay

Cell viability was determined using the CellTiter MTS Assay (Promega) according to the manufacturer’s protocol. CellTiter solution was added to complete medium to a final concentration of 1x and incubated with the samples for one hour at 37LJ. After, 100μL from each sample was transferred to a 96-well plate. The absorbance at 492nm was measured using a plate reader. After subtracting the absorbance reading of cell-free MTS solution, the values were normalized to percent of the control wells.

### Heme Assay

Heme levels in the cells were determined as previously described.(72) Briefly, cell lysates were collected from ∼80% confluent 6-well plates following incubation with RIPA buffer. 2M oxalic acid solution was prepared and 10 μL of the cell lysate was added to 200 μL oxalic acid in a 1.5mL protein lo-bind tube. Samples were heated to 95 °C for 30 min, followed by centrifugation for 10 min at 1000× g at 4 °C to remove the debris. 100 μL of the supernatant was transferred to a 96-well plate, and the fluorescence was assessed at 405/600 nm by using a plate reader.

### RPA Assay

Free mitochondrial iron was measured using RPA according to the manufacturer’s protocol (Squarix Biotechnology GmbH, Marl, Germany). Cells were cultured in a glass-bottom 48-well plate and treated overnight with 10μM CoPP. Cells were washed with Hank’s Buffered Saline Solution (HBSS), then incubated with 5μM RPA for 20 minutes at 37 °C. Cells were washed twice with HBSS and incubated for an additional 15 min in a dye-free buffer at 37 °C. RPA fluorescence was determined by confocal imaging using a Leica SP8 microscope.

### Raman Spectroscopy

Raman spectroscopy was performed to determine how glucose was metabolized in the collectively migrating cells. Adapting the protocol from Bagheri *et al.* (106), 12 mm coverslips were sterilized in 100% ethanol for one hour and dried with lint-free paper. Coverslips were then incubated under UV for 20 minutes to complete sterilization, moved into a 24-well plate and functionalized using a plasma cleaner. A 50µL 6 mg/mL Collagen I gel seeded with 6250 MDA-MB-231 WT cells was seeded on top of each coverslip. Media was changed every 2 days for the first 5 days. On the 5th day of culture, collagen gels were switched to a starvation media for 6 hours containing DMEM without Glucose, FBS or Sodium Pyruvate (Gibco #11966-025) and then switched into normal DMEM containing 4.5 mg/mL D-Glucose (DLM-2062, Cambridge Isotope Laboratories) + 1mM Sodium Pyruvate (Catalog #: 25-000-CI-1) + 10% FBS for 36 hours. Collagen I gels were washed with PBS and fixed for 20 minutes at room temperature in 4% PFA solution. This was followed by 3x washes in PBS and samples were stored at 4°C until mounting onto slides.

The SRS signal from the cell samples were collected from an upright laser-scanning microscope (DIY multiphoton, Olympus), which was equipped with a 25x water objective (XLPLN, WMP2, 1.05 NA, Olympus). The synchronized pulsed pump beam (tunable 720–990 nm wavelength, 5– 6 ps pulse width, and 80 MHz repetition rate) and stokes beam (wavelength at 1031nm, 6 ps pulse width, and 80 MHz repetition rate) from a picoEmerald system (Applied Physics & Electronics) were coupled and introduced into the microscope. The transmitted signal from the pump and Stokes beams after interacting with the samples was collected by a high NA oil condenser (1.4 NA).

A shortpass filter (950 nm, Thorlabs) was used to completely block the Stokes beam and transmit the pump beam onto a Si photodiode array for detecting the stimulated Raman loss signal. The output current from the photodiode array was terminated, filtered, and demodulated by a lock-in amplifier at 20 MHz. The demodulated signal was fed into the FV3000 software module FV-OSR (Olympus) to form images using laser scanning. The SRS-lipid images were obtained at vibrational mode of 2850 cm^-1^ in a single frame of 512 x 512 pixels, at a dwell time 80 μs. The C-D vibrational modes of deuterium labeled lipids derived D-Glucose labeling were acquired at Raman peak 2143 cm^-1^ in a single frame of 512 x 512 pixels, at a dwell time 80 μs. A background signal was acquired at 2190 cm^-1^ in the Raman cell silent region with the same parameters and subtracted from all SRS images acquired in the same region of interest using ImageJ. To quantify the lipid turnover rate, the intensity of C-D lipid vibrational modes (peak intensity at 2140 cm-1) was divided by the intensity of C-H vibrational modes (peak intensity at 2850 cm-1), resulting in CD/CH ratiometric images (107). Minor intensity adjustment (brightness and/or contrast) was performed in ImageJ. 9 regions of interest from 3 independent experiments were imaged and analyzed, the results were further quantified in ImageJ and plotted in GraphPad 10.

The two-photon fluorescent microscopy was used to collect the NADH and FAD signal from the samples. The NADH signal was excited by 780 nm, the emission wavelength was collected at 460 nm. The FAD signal was excited by 860 nm, the emission wavelength was collected at 515 nm. The ratiometric images were determined by NADH/FAD. 9 regions of interest from 3 independent experiments were imaged and analyzed. The ratio was further quantified in ImageJ and plotted in GraphPad 10.

## Acknowledgements

Support for this study was provided by National Science Foundation CAREER award MCB-1651855 (S.I.F.), National Science Foundation grant DMS-1953469, National Cancer Institute (NCI) grant CA274502, American Cancer Society Research Scholar Grant RSG-21-033-01-CSM (S.I.F.). This work was also supported by grants from the NCI (RO1CA262794, R01CA174869, R01CA268179, and R01CA236386) to J.Y. A.M.F. is supported by the TRDRP Postdoctoral Award T32FT4922. Z.H. is supported by the Pfizer-Cell Signaling San Diego Fellowship. S.R. is supported by the NCI CT2 Postdoctoral Training Grant (2T32CA121938). We would like to thank the UC San Diego School of Medicine Microscopy Core, which is supported by the National Institute of Neurological Disorders and Stroke grant P30NS047101.

## Author Contributions

W.D.L. designed experiments, conducted experimentation, analyzed data, and contributed to the writing of the manuscript. S.I.F. conceived of the project, assisted in experimental design and interpretation, and contributed to the writing of the manuscript. A.M.F., Z.H., and J.Y. generated the PDO model and S.R. performed the PDO experiments. A.W. performed the shACO1 western blots, and M.Z.R. imaged the cytoskeletal inhibitor gels, Rac1 stains, heme assays, and mitochondrial iron assays.

---

**Figure S1:**
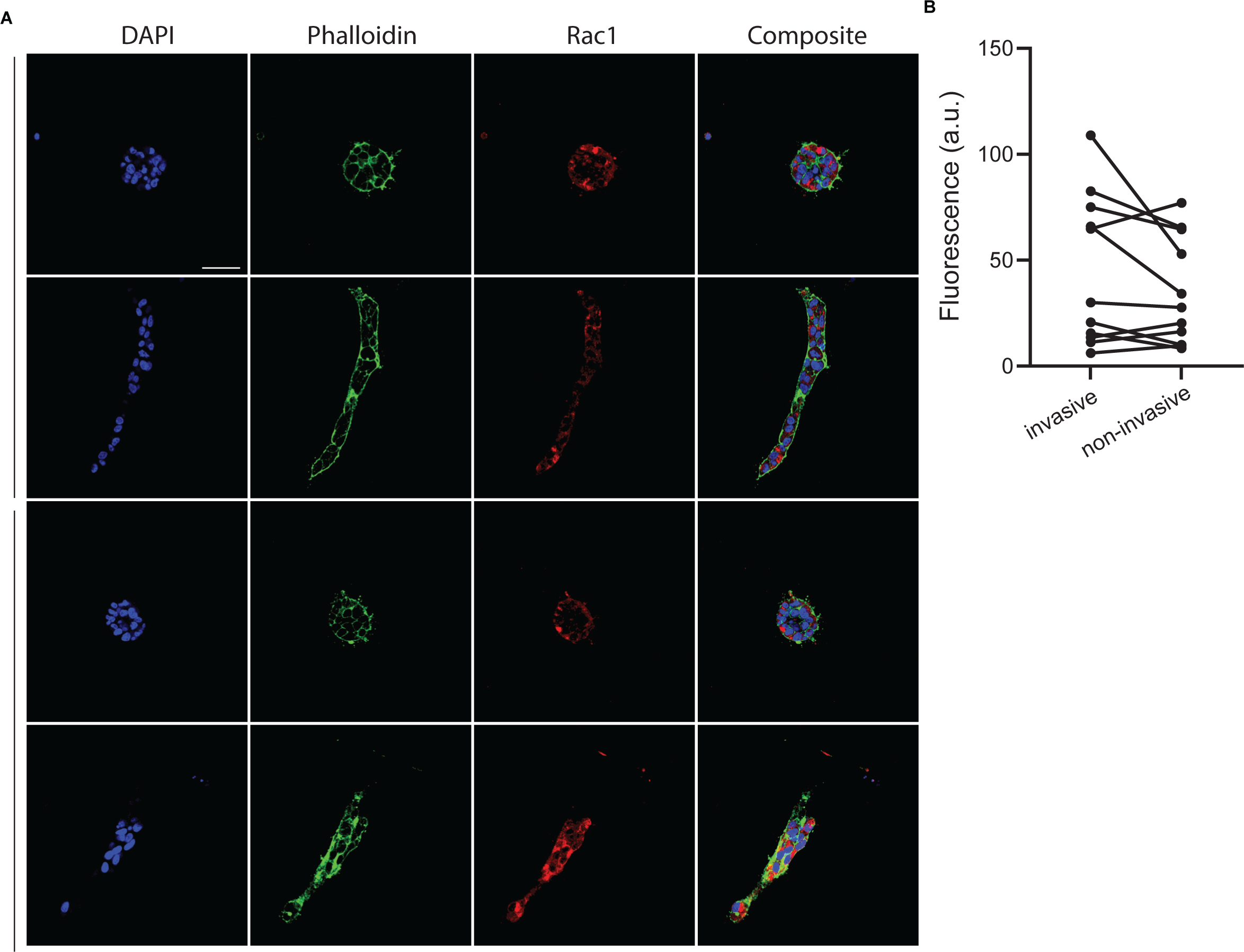
Immunostaining of Rac1 shows no quantitative differences between invasive and non-invasive phenotypes. MDA-MB-231 cells cultured in 6mg/ml Col1 for seven days were fixed and stained to show Rac1/2 expression levels. (A) Representative micrographs from confocal microscopy show the nuclei, actin, and Rac1 protein levels. The average fluorescence intensity was measured within each multicellular structure. Scale bar = 200µm. (B) Pairwise comparisons of structures from within the same collagen gel were used to account for fluctuations in fluorescence intensity arising from staining or imaging factors.

**Figure S2:**
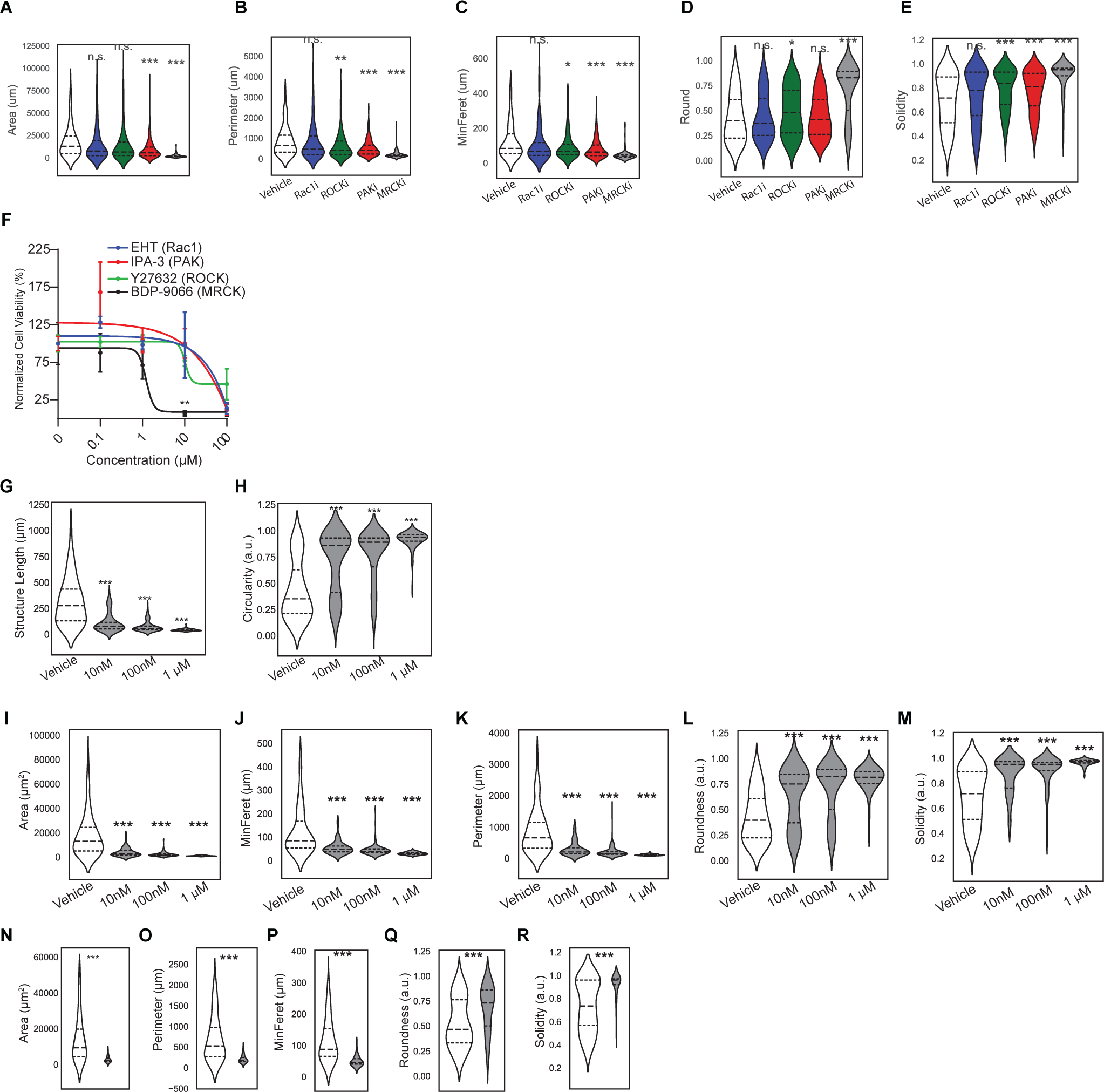
Effects of cytoskeletal inhibitors on collective migration phenotypes and viability. **(A-E)** Shape descriptors of MDAs embedded in HD Col1 for seven days after initial 24hr treatment with cytoskeletal inhibitors. **(F)** MTS assay of MDAs embedded in HD Col1 for seven days after initial 24hr treatment with cytoskeletal inhibitors. **(G-M)** Shape descriptors of MDAs embedded in HD Col1 for seven days after initial 24hr treatment with different concentrations of MRCK inhibitors. **(N-R)** Shape descriptors of 4T1s embedded in HD Col1 for seven days after initial 24hr treatment with MRCK inhibitor BDP-9066 at 1uM.

**Figure S3:**
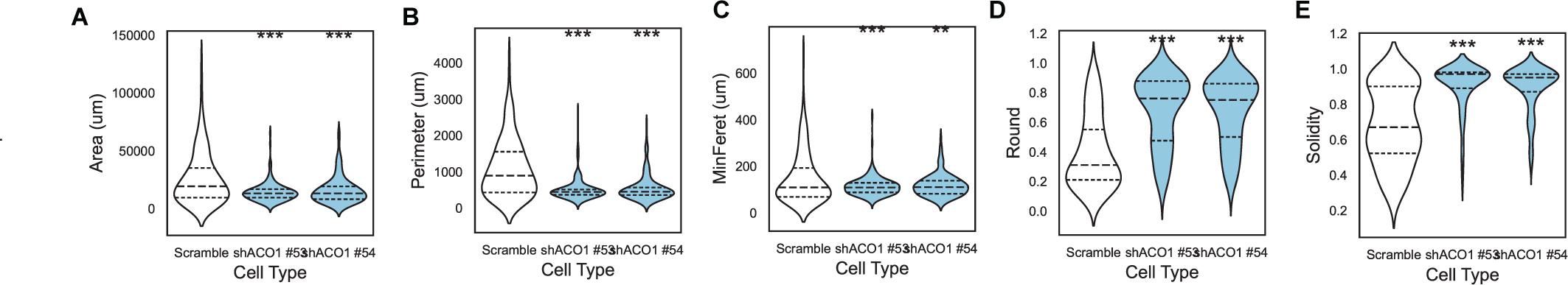
shACO1 shape quantification. (A) WesternShape descriptors of shACO1 MDAs embedded in HD Col1 for seven days.

**Figure S4:**
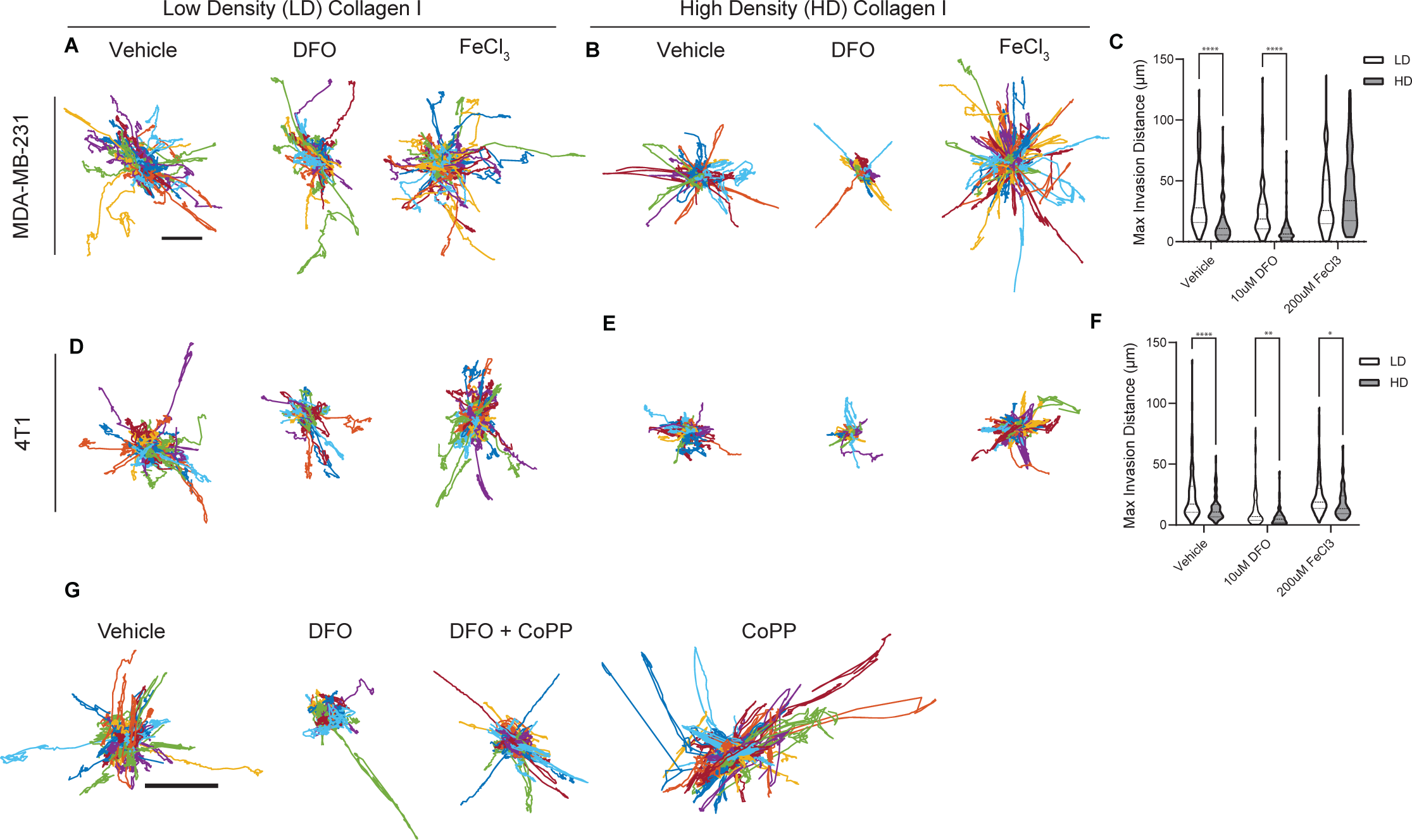
MDA single cell migration trajectories in HD Col1. **(A-C)** Trajectories of MDAs embedded in LD (A) or HD (B) matrices and the associated max invasion distances (C). **(D-F)** Trajectories of MDAs embedded in LD (D) or HD (E) matrices and the associated max invasion distances (F). **(G)** Trajectories of MDAs in HD matrices treated with 0.1% DMSO, 10uM DFO, 10uM DFO + 100uM CoPP, or 100uM CoPP.

**Figure S5:**
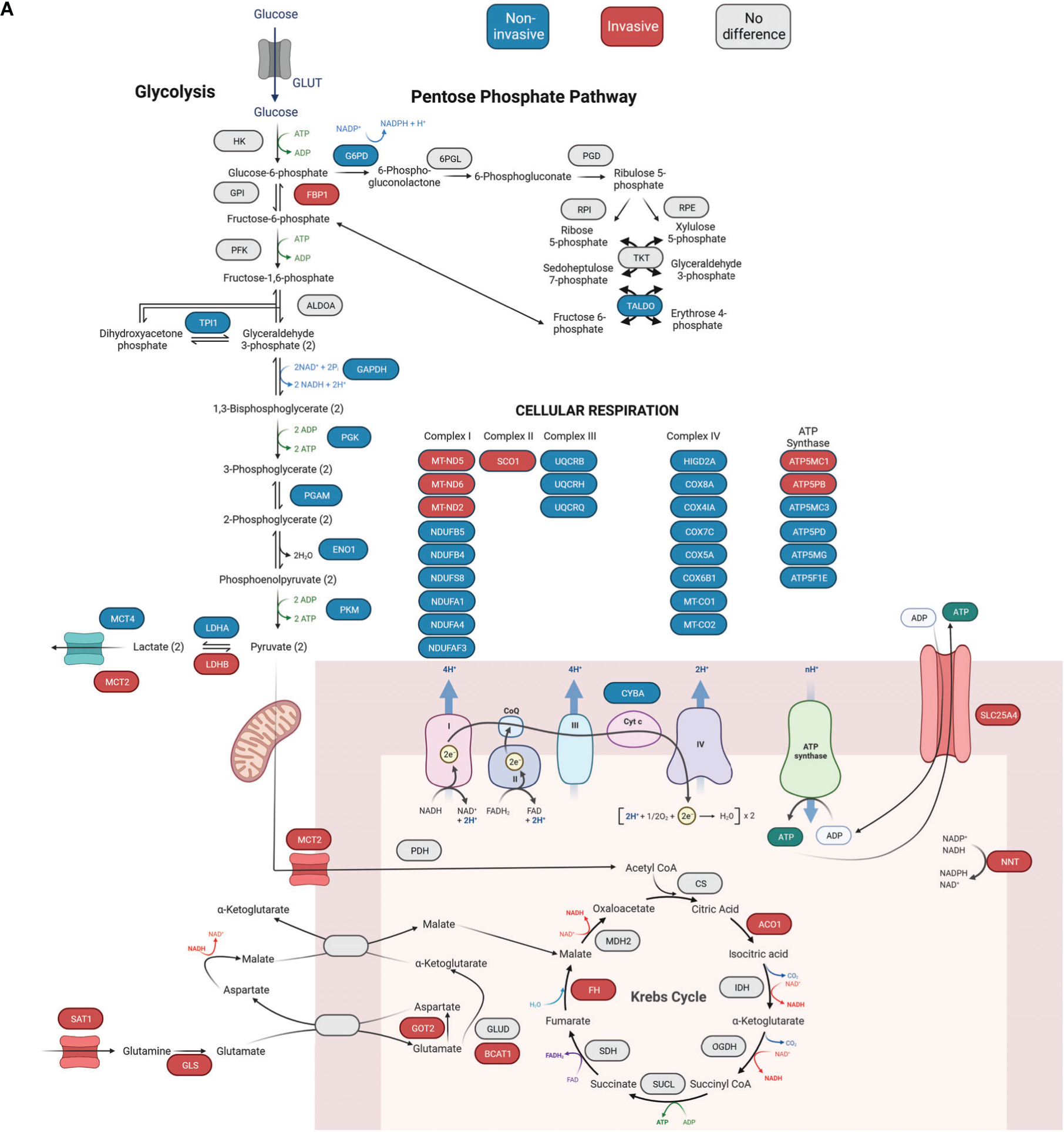
Differentially expressed genes mapped to glucose metabolism. (A) Essential proteins involved in glucose metabolism through glycolysis, the TCA cycle, and oxidative phosphorylation. Shunting from glycolysis to the pentose phosphate pathway is also shown, as are anapleirotic sources of the TCA cycle.

**Figure S6:**
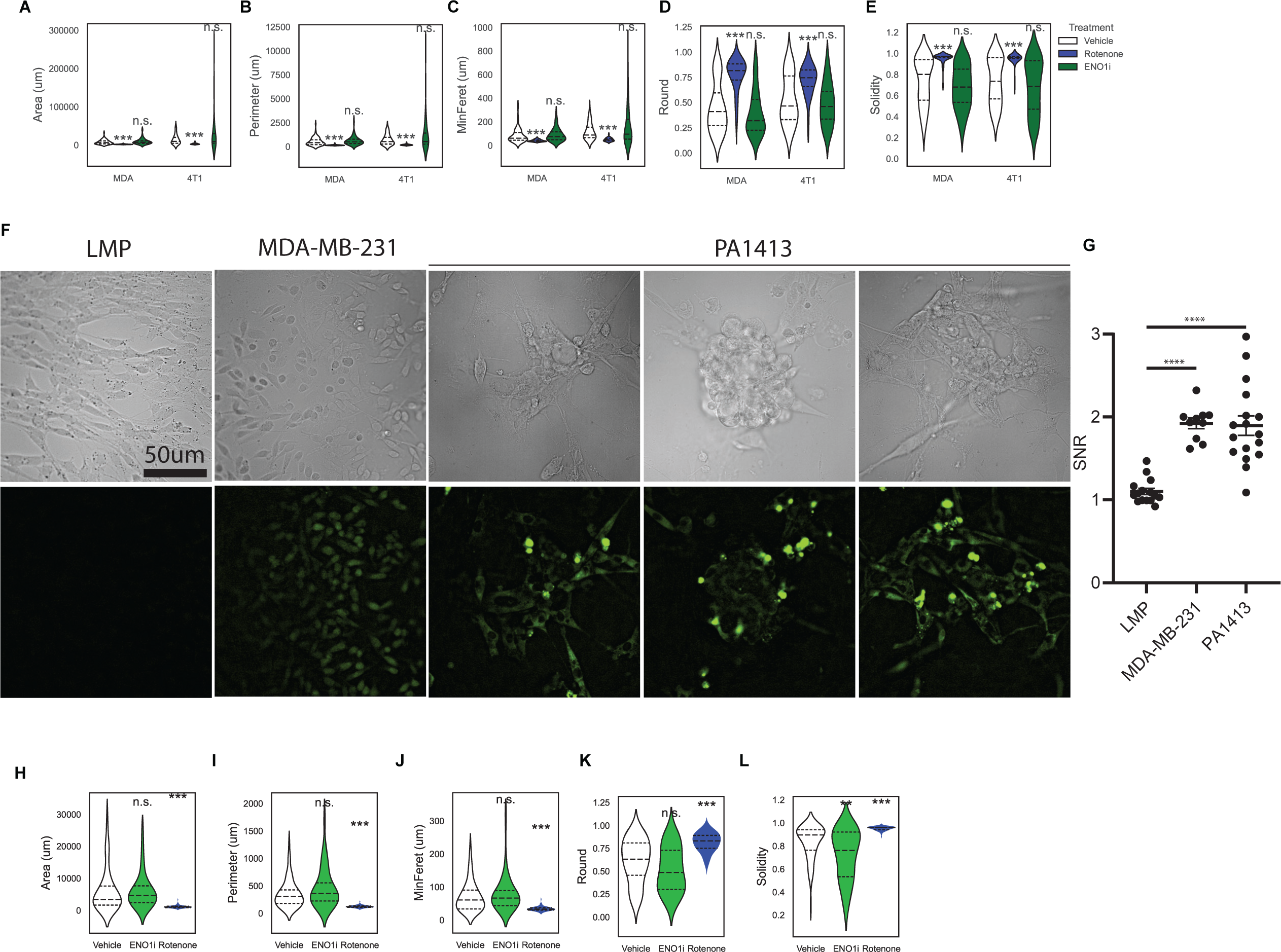
Metabolic inhibitors extended shape descriptors and PDO model validation. **(A-E)** Shape descriptors of MDAs and 4T1s embedded in HD Col1 for seven days after initial 24hr treatment with inhibitors targeting Complex I of the electron transport chain and glycolytic enzyme ENO1. **(F)** Cells from the PDOs after excision from mouse xenografts were validated to be human via staining with a human-specific antibody. While mouse LMP cells have no signal, the PA1413 model shows staining at a similar level as MDAs. **(G)** Quantification of the signal-to-noise ratio (SNR). **(H-L)** Shape descriptors of the PA1413 PDO model after treatment with inhibitors targeting Complex I of the electron transport chain and glycolytic enzyme ENO1.

**Figure S7:**
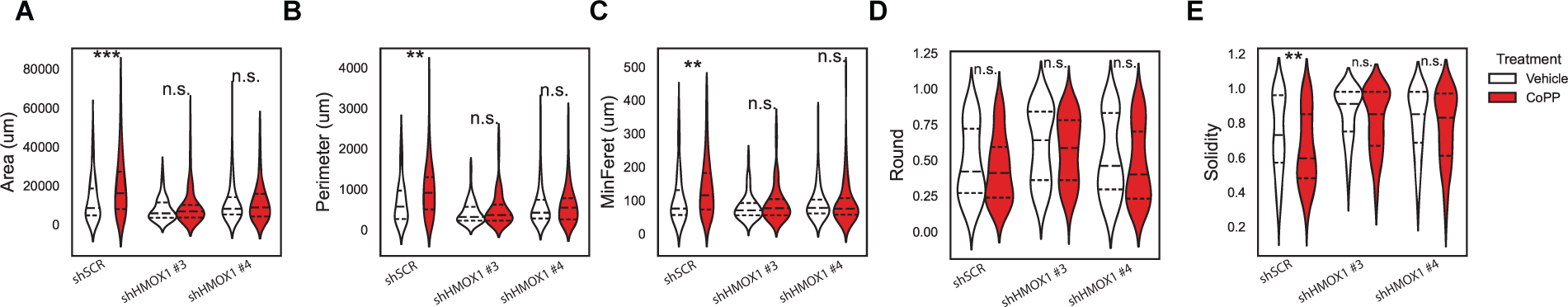
Metabolic inhibitors extended shape descriptors and PDO model validation. **(A-E)** Shape descriptors of MDAs embedded in HD Col1 for seven days with media supplemented with 10uM CoPP.

Table S1 - scRNA-sequencing results.

Table S2 - Rho GTPase and actin-regulating DE genes

Table S3 - Iron DE genes

Table S4 - Metabolism DE genes

Table S5 - Model Parameters and Reactions

## References

1. H. Dillekås, M. S. Rogers, O. Straume, Are 90% of deaths from cancer caused by metastases? Cancer Med. 8, 5574–5576 (2019).

2. A. Wells, J. Grahovac, S. Wheeler, B. Ma, D. Lauffenburger, Targeting tumor cell motility as a strategy against invasion and metastasis. Trends Pharmacol. Sci. 34, 283–289 (2013).

3. T. D. Palmer, W. J. Ashby, J. D. Lewis, A. Zijlstra, Targeting tumor cell motility to prevent metastasis. Adv. Drug Deliv. Rev. 63, 568–581 (2011).

4. K. Ganesh, J. Massagué, Targeting metastatic cancer. Nat. Med. 27, 34–44 (2021).

5. S. A. Patel, P. Rodrigues, L. Wesolowski, S. Vanharanta, Genomic control of metastasis. Br. J. Cancer 124, 3–12 (2021).

6. P. Priestley, et al., Pan-cancer whole-genome analyses of metastatic solid tumours. Nature 575, 210–216 (2019).

7. S. Turajlic, C. Swanton, Metastasis as an evolutionary process. Science 352, 169–175 (2016).

8. B. Vogelstein, et al., Cancer genome landscapes. Science 339, 1546–1558 (2013).

9. P. Rodrigues, et al., NF-κB--Dependent Lymphoid Enhancer Co-option Promotes Renal Carcinoma MetastasisCross-Lineage Enhancer Co-option Promotes RCC Metastasis. Cancer Discov. 8, 850–865 (2018).

10. K. Ganesh, et al., Author Correction: L1CAM defines the regenerative origin of metastasis-initiating cells in colorectal cancer. Nat Cancer 1, 1128 (2020).

11. R. Eyre, et al., Microenvironmental IL1β promotes breast cancer metastatic colonisation in the bone via activation of Wnt signalling. Nat. Commun. 10, 5016 (2019).

12. N. Mohammadi Ghahhari, et al., Cooperative interaction between ERα and the EMT-inducer ZEB1 reprograms breast cancer cells for bone metastasis. Nat. Commun. 13, 2104 (2022).

13. A. A. Chakraborty, et al., Histone demethylase KDM6A directly senses oxygen to control chromatin and cell fate. Science 363, 1217–1222 (2019).

14. C. Wiel, et al., BACH1 Stabilization by Antioxidants Stimulates Lung Cancer Metastasis. Cell 178, 330–345.e22 (2019).

15. D. A. Lawson, K. Kessenbrock, R. T. Davis, N. Pervolarakis, Z. Werb, Tumour heterogeneity and metastasis at single-cell resolution. Nat. Cell Biol. 20, 1349–1360 (2018).

16. K. J. Cheung, A. J. Ewald, A collective route to metastasis: Seeding by tumor cell clusters. Science 352, 167–169 (2016).

17. P. Friedl, D. Gilmour, Collective cell migration in morphogenesis, regeneration and cancer. Nat. Rev. Mol. Cell Biol. 10, 445–457 (2009).

18. K. J. Cheung, E. Gabrielson, Z. Werb, A. J. Ewald, Collective invasion in breast cancer requires a conserved basal epithelial program. Cell 155, 1639–1651 (2013).

19. N. Aceto, et al., Circulating tumor cell clusters are oligoclonal precursors of breast cancer metastasis. Cell 158, 1110–1122 (2014).

20. A. Haeger, M. Krause, K. Wolf, P. Friedl, Cell jamming: collective invasion of mesenchymal tumor cells imposed by tissue confinement. Biochim. Biophys. Acta 1840, 2386–2395 (2014).

21. D. O. Velez, et al., 3D collagen architecture induces a conserved migratory and transcriptional response linked to vasculogenic mimicry. Nat. Commun. 8, 1651 (2017).

22. S. K. Ranamukhaarachchi, et al., Macromolecular crowding tunes 3D collagen architecture and cell morphogenesis. Biomater Sci 7, 618–633 (2019).

23. N. F. Boyd, et al., Mammographic breast density as an intermediate phenotype for breast cancer. Lancet Oncol. 6, 798–808 (2005).

24. N. F. Boyd, et al., Mammographic density and the risk and detection of breast cancer. N. Engl. J. Med. 356, 227–236 (2007).

25. J. J. Northey, et al., Stiff stroma increases breast cancer risk by inducing the oncogene ZNF217. J. Clin. Invest. 130, 5721–5737 (2020).

26. M. J. Paszek, et al., Tensional homeostasis and the malignant phenotype. Cancer Cell 8, 241–254 (2005).

27. D. O. Velez, et al., 3D collagen architecture regulates cell adhesion through degradability, thereby controlling metabolic and oxidative stress. Integr. Biol. 11, 221–234 (2019).

28. K. Chen, et al., Phenotypically supervised single-cell sequencing parses within-cell-type heterogeneity. iScience 24, 101991 (2021).

29. K. Chen, et al., A phenotypically supervised single-cell analysis protocol to study within-cell-type heterogeneity of cultured mammalian cells. STAR Protoc 2, 100561 (2021).

30. C. Dessimoz, N. Škunca, The gene ontology handbook (Springer Nature, 2017).

31. A. Tomczak, et al., Interpretation of biological experiments changes with evolution of the Gene Ontology and its annotations. Sci. Rep. 8, 5115 (2018).

32. L. Dong, et al., DIAPH3 promoted the growth, migration and metastasis of hepatocellular carcinoma cells by activating beta-catenin/TCF signaling. Mol. Cell. Biochem. 438, 183– 190 (2018).

33. R. S. Maul, et al., EPLIN regulates actin dynamics by cross-linking and stabilizing filaments. J. Cell Biol. 160, 399–407 (2003).

34. H. Phee, et al., Pak2 is required for actin cytoskeleton remodeling, TCR signaling, and normal thymocyte development and maturation. Elife 3, e02270 (2014).

35. S. Liu, The ROCK signaling and breast cancer metastasis. Mol. Biol. Rep. 38, 1363–1366 (2011).

36. J. L. Ouderkirk, M. Krendel, Non-muscle myosins in tumor progression, cancer cell invasion, and metastasis. Cytoskeleton 71, 447–463 (2014).

37. K. A. Makowska, R. E. Hughes, K. J. White, C. M. Wells, M. Peckham, Specific Myosins Control Actin Organization, Cell Morphology, and Migration in Prostate Cancer Cells. Cell Rep. 13, 2118–2125 (2015).

38. C. Lei, et al., miR-143 and miR-145 inhibit gastric cancer cell migration and metastasis by suppressing MYO6. Cell Death Dis. 8, e3101 (2017).

39. T. Omelchenko, Regulation of collective cell migration by RhoGAP myosin IXA. Small GTPases 3, 213–218 (2012).

40. F.-S. Yi, et al., TSAd Plays a Major Role in Myo9b-Mediated Suppression of Malignant Pleural Effusion by Regulating TH1/TH17 Cell Response. J. Immunol. 205, 2926–2935 (2020).

41. E. Nishida, S. Maekawa, H. Sakai, Cofilin, a protein in porcine brain that binds to actin filaments and inhibits their interactions with myosin and tropomyosin. Biochemistry 23, 5307–5313 (1984).

42. P. Scherle, T. Behrens, L. M. Staudt, Ly-GDI, a GDP-dissociation inhibitor of the RhoA GTP-binding protein, is expressed preferentially in lymphocytes. Proc. Natl. Acad. Sci. U. S. A. 90, 7568–7572 (1993).

43. C. N. Adra, et al., Identification of a novel protein with GDP dissociation inhibitor activity for the ras-like proteins CDC42Hs and rac I. Genes Chromosomes Cancer 8, 253–261 (1993).

44. R. Cmejla, et al., Human MRCKalpha is regulated by cellular iron levels and interferes with transferrin iron uptake. Biochem. Biophys. Res. Commun. 395, 163–167 (2010).

45. P. Y. He, et al., Inhibition of cell migration and invasion by miRLJ29aLJ3p in a colorectal cancer cell line through suppression of CDC42BPA mRNA expression. Oncol. Rep. 38, 3554–3566 (2017).

46. J. I. Lachowicz, et al., Thymosin β4 Is an Endogenous Iron Chelator and Molecular Switcher of Ferroptosis. Int. J. Mol. Sci. 23 (2022).

47. K. A. O’Donnell, et al., Activation of transferrin receptor 1 by c-Myc enhances cellular proliferation and tumorigenesis. Mol. Cell. Biol. 26, 2373–2386 (2006).

48. T. A. Rouault, The role of iron regulatory proteins in mammalian iron homeostasis and disease. Nat. Chem. Biol. 2, 406–414 (2006).

49. B. Győrffy, Survival analysis across the entire transcriptome identifies biomarkers with the highest prognostic power in breast cancer. Comput. Struct. Biotechnol. J. 19, 4101–4109 (2021).

50. G. F. Vile, R. M. Tyrrell, Oxidative stress resulting from ultraviolet A irradiation of human skin fibroblasts leads to a heme oxygenase-dependent increase in ferritin. J. Biol. Chem. 268, 14678–14681 (1993).

51. R. S. Eisenstein, D. Garcia-Mayol, W. Pettingell, H. N. Munro, Regulation of ferritin and heme oxygenase synthesis in rat fibroblasts by different forms of iron. Proc. Natl. Acad. Sci. U. S. A. 88, 688–692 (1991).

52. G. F. Vile, S. Basu-Modak, C. Waltner, R. M. Tyrrell, Heme oxygenase 1 mediates an adaptive response to oxidative stress in human skin fibroblasts. Proc. Natl. Acad. Sci. U. S. A. 91, 2607–2610 (1994).

53. P. N. Dayani, M. C. Bishop, K. Black, P. M. Zeltzer, Desferoxamine (DFO)--mediated iron chelation: rationale for a novel approach to therapy for brain cancer. J. Neurooncol. 67, 367–377 (2004).

54. D. M. Ward, S. M. Cloonan, Mitochondrial Iron in Human Health and Disease. Annu. Rev. Physiol. 81, 453–482 (2019).

55. G. Liu, et al., Heme biosynthesis depends on previously unrecognized acquisition of iron-sulfur cofactors in human amino-levulinic acid dehydratase. Nat. Commun. 11, 6310 (2020).

56. F. Petrat, et al., Selective determination of mitochondrial chelatable iron in viable cells with a new fluorescent sensor. Biochem. J 362, 137–147 (2002).

57. S.-I. Ogura, et al., The effect of 5-aminolevulinic acid on cytochrome c oxidase activity in mouse liver. BMC Res. Notes 4, 66 (2011).

58. Y. Sugiyama, et al., The heme precursor 5-aminolevulinic acid disrupts the Warburg effect in tumor cells and induces caspase-dependent apoptosis. Oncol. Rep. 31, 1282–1286 (2014).

59. S. Sohoni, et al., Elevated Heme Synthesis and Uptake Underpin Intensified Oxidative Metabolism and Tumorigenic Functions in Non-Small Cell Lung Cancer Cells. Cancer Res. 79, 2511–2525 (2019).

60. H. J. Kim, O. Khalimonchuk, P. M. Smith, D. R. Winge, Structure, function, and assembly of heme centers in mitochondrial respiratory complexes. Biochim. Biophys. Acta 1823, 1604–1616 (2012).

61. A. Signes, E. Fernandez-Vizarra, Assembly of mammalian oxidative phosphorylation complexes I-V and supercomplexes. Essays Biochem. 62, 255–270 (2018).

62. K. Urbańska, A. Orzechowski, Unappreciated Role of LDHA and LDHB to Control Apoptosis and Autophagy in Tumor Cells. Int. J. Mol. Sci. 20 (2019).

63. T. Zhang, et al., ENO1 suppresses cancer cell ferroptosis by degrading the mRNA of iron regulatory protein 1. Nat Cancer 3, 75–89 (2022).

64. Q. Liu, et al., circ-Pank1 promotes dopaminergic neuron neurodegeneration through modulating miR-7a-5p/α-syn pathway in Parkinson’s disease. Cell Death Dis. 13, 477 (2022).

65. F. Wang, et al., ALCAM regulates multiple myeloma chemoresistant side population. Cell Death Dis. 13, 136 (2022).

66. D.-W. Jung, et al., A unique small molecule inhibitor of enolase clarifies its role in fundamental biological processes. ACS Chem. Biol. 8, 1271–1282 (2013).

67. M. Morales, X. Xue, Targeting iron metabolism in cancer therapy. Theranostics 11, 8412– 8429 (2021).

68. D. Basuli, et al., Iron addiction: a novel therapeutic target in ovarian cancer. Oncogene 36, 4089–4099 (2017).

69. P. Lelièvre, L. Sancey, J.-L. Coll, A. Deniaud, B. Busser, Iron Dysregulation in Human Cancer: Altered Metabolism, Biomarkers for Diagnosis, Prognosis, Monitoring and Rationale for Therapy. Cancers 12 (2020).

70. K. Bajbouj, J. Shafarin, M. Hamad, High-Dose Deferoxamine Treatment Disrupts Intracellular Iron Homeostasis, Reduces Growth, and Induces Apoptosis in Metastatic and Nonmetastatic Breast Cancer Cell Lines. Technol. Cancer Res. Treat. 17, 1533033818764470 (2018).

71. C. Chen, P. Liu, X. Duan, M. Cheng, L. X. Xu, Deferoxamine-induced high expression of TfR1 and DMT1 enhanced iron uptake in triple-negative breast cancer cells by activating IL-6/PI3K/AKT pathway. Onco. Targets. Ther. 12, 4359–4377 (2019).

72. C. Chen, S. Wang, P. Liu, Deferoxamine Enhanced Mitochondrial Iron Accumulation and Promoted Cell Migration in Triple-Negative MDA-MB-231 Breast Cancer Cells Via a ROS-Dependent Mechanism. Int. J. Mol. Sci. 20 (2019).

73. C. Zapp, et al., Mechanoradicals in tensed tendon collagen as a source of oxidative stress. Nat. Commun. 11, 2315 (2020).

74. M. J. Harrington, A. Masic, N. Holten-Andersen, J. H. Waite, P. Fratzl, Iron-clad fibers: a metal-based biological strategy for hard flexible coatings. Science 328, 216–220 (2010).

75. B. Wegiel, Z. Nemeth, M. Correa-Costa, A. C. Bulmer, L. E. Otterbein, Heme oxygenase-1: a metabolic nike. Antioxid. Redox Signal. 20, 1709–1722 (2014).

76. A. Jozkowicz, H. Was, J. Dulak, Heme oxygenase-1 in tumors: is it a false friend? Antioxid. Redox Signal. 9, 2099–2117 (2007).

77. K. N. Luu Hoang, J. E. Anstee, J. N. Arnold, The Diverse Roles of Heme Oxygenase-1 in Tumor Progression. Front. Immunol. 12, 658315 (2021).

78. H. Was, J. Dulak, A. Jozkowicz, Heme oxygenase-1 in tumor biology and therapy. Curr. Drug Targets 11, 1551–1570 (2010).

79. Q. Li, et al., Heme Oxygenase-1 Inhibits Tumor Metastasis Mediated by Notch1 Pathway in Murine Mammary Carcinoma. Oncol. Res. 27, 643–651 (2019).

80. J.-R. Tsai, et al., High expression of heme oxygenase-1 is associated with tumor invasiveness and poor clinical outcome in non-small cell lung cancer patients. Cell. Oncol. 35, 461–471 (2012).

81. N. A. Gandini, et al., Heme Oxygenase-1 Has an Antitumor Role in Breast Cancer. Antioxid. Redox Signal. 30, 2030–2049 (2019).

82. M. Mascaró, et al., Nuclear Localization of Heme Oxygenase-1 in Pathophysiological Conditions: Does It Explain the Dual Role in Cancer? Antioxidants (Basel) 10 (2021).

83. L. D. Miller, et al., An iron regulatory gene signature predicts outcome in breast cancer. Cancer Res. 71, 6728–6737 (2011).

84. J. V. Dietz, J. L. Fox, O. Khalimonchuk, Down the Iron Path: Mitochondrial Iron Homeostasis and Beyond. Cells 10 (2021).

85. M. Doss, F. Sixel-Dietrich, F. Verspohl, “Glucose effect” and rate limiting function of uroporphyrinogen synthase on porphyrin metabolism in hepatocyte culture: relationship with human acute hepatic porphyrias. J. Clin. Chem. Clin. Biochem. 23, 505–513 (1985).

86. P. Kaur, et al., Activated heme synthesis regulates glycolysis and oxidative metabolism in breast and ovarian cancer cells. PLoS One 16, e0260400 (2021).

87. P. Kaur, et al., Increased heme synthesis in ovarian cancer and triple negative breast cancer cells leads to downregulation of glycolysis, mitochondrial respiration, and cell migration. FASEB J. 34, 1–1 (2020).

88. X. G. Zhu, et al., Functional Genomics In Vivo Reveal Metabolic Dependencies of Pancreatic Cancer Cells. Cell Metab. 33, 211–221.e6 (2021).

89. M. Miyazawa, A. R. Bogdan, Y. Tsuji, Perturbation of Iron Metabolism by Cisplatin through Inhibition of Iron Regulatory Protein 2. Cell Chem Biol 26, 85–97.e4 (2019).

90. C. K. Huang, Y. Sun, L. Lv, Y. Ping, ENO1 and Cancer. Mol Ther Oncolytics 24, 288–298 (2022).

91. T. M. Ashton, W. G. McKenna, L. A. Kunz-Schughart, G. S. Higgins, Oxidative Phosphorylation as an Emerging Target in Cancer Therapy. Clin. Cancer Res. 24, 2482– 2490 (2018).

92. P. E. Porporato, et al., A mitochondrial switch promotes tumor metastasis. Cell Rep. 8, 754– 766 (2014).

93. X. L. Zu, M. Guppy, Cancer metabolism: facts, fantasy, and fiction. Biochem. Biophys. Res. Commun. 313, 459–465 (2004).

94. K. Karrobi, et al., Fluorescence Lifetime Imaging Microscopy (FLIM) reveals spatial-metabolic changes in 3D breast cancer spheroids. Sci. Rep. 13, 3624 (2023).

95. D. F. Quail, J. A. Joyce, Microenvironmental regulation of tumor progression and metastasis. Nat. Med. 19, 1423–1437 (2013).

96. J. A. Kenkel, et al., An Immunosuppressive Dendritic Cell Subset Accumulates at Secondary Sites and Promotes Metastasis in Pancreatic Cancer. Cancer Res. 77, 4158–4170 (2017).

97. W. W. Tseng, et al., Development of an orthotopic model of invasive pancreatic cancer in an immunocompetent murine host. Clin. Cancer Res. 16, 3684–3695 (2010).

98. J. Schindelin, et al., Fiji: an open-source platform for biological-image analysis. Nat. Methods 9, 676–682 (2012).

99. S. I. Fraley, et al., Three-dimensional matrix fiber alignment modulates cell migration and MT1-MMP utility by spatially and temporally directing protrusions. Sci. Rep. 5, 14580 (2015).

100. E. Y. Chen, et al., Enrichr: interactive and collaborative HTML5 gene list enrichment analysis tool. BMC Bioinformatics 14, 128 (2013).

101. M. V. Kuleshov, et al., Enrichr: a comprehensive gene set enrichment analysis web server 2016 update. Nucleic Acids Res. 44, W90–7 (2016).

102. Z. Xie, et al., Gene Set Knowledge Discovery with Enrichr. Curr Protoc 1, e90 (2021).

103. M. Uhlén, et al., Proteomics. Tissue-based map of the human proteome. Science 347, 1260419 (2015).

104. R. T. Powell, et al., Pharmacologic profiling of patient-derived xenograft models of primary treatment-naïve triple-negative breast cancer. Sci. Rep. 10, 17899 (2020).

105. K. P. Guillen, et al., A human breast cancer-derived xenograft and organoid platform for drug discovery and precision oncology. Nat Cancer 3, 232–250 (2022).

106. P. Bagheri, et al., Bioorthogonal Chemical Imaging of Cell Metabolism Regulated by Aromatic Amino Acids. J. Vis. Exp. (2023) 10.3791/65121.

107. L. Zhang, et al., Spectral tracing of deuterium for imaging glucose metabolism. Nat Biomed Eng 3, 402–413 (2019).

108. A. Lawen, D. J. R. Lane, Mammalian iron homeostasis in health and disease: uptake, storage, transport, and molecular mechanisms of action. Antioxid. Redox Signal. 18, 2473– 2507 (2013).

109. C. P. Anderson, M. Shen, R. S. Eisenstein, E. A. Leibold, Mammalian iron metabolism and its control by iron regulatory proteins. Biochim. Biophys. Acta 1823, 1468–1483 (2012).

